# TDP-43 and HSP70 phase separate into anisotropic, intranuclear liquid spherical annuli

**DOI:** 10.1101/2020.03.28.985986

**Authors:** Haiyang Yu, Shan Lu, Kelsey Gasior, Digvijay Singh, Olga Tapia, Sonia Vazquez-Sanchez, Divek Toprani, Melinda S. Beccari, John R. Yates, Sandrine Da Cruz, Jay M. Newby, Miguel Larfaga, Amy S. Gladfelter, Elizabeth Villa, Don W. Cleveland

**Author notes:** Florida State University Department of Mathematics, 1017 Academic Way, Tallahassee, FL 32304. These authors contribute equally to this work (Shan Lu, Kelsey Gasior).

## Abstract

The RNA binding protein TDP-43 naturally phase separates within cell nuclei and forms cytoplasmic aggregates in age-related neurodegenerative diseases. Here we show that acetylation-mediated inhibition of TDP-43 binding to RNA produces co-de-mixing of acetylated and unmodified TDP-43 into symmetrical, intranuclear spherical annuli whose shells and cores have liquid properties. Shells are anisotropic, like liquid crystals. Consistent with our modelling predictions that annulus formation is driven by components with strong self-interactions but weak interaction with TDP-43, the major components of annuli cores are identified to be HSP70 family proteins, whose chaperone activity is required to maintain liquidity of the core. Proteasome inhibition, mimicking reduction in proteasome activity during aging, induces TDP-43-containing annuli in neurons in rodents. Thus, we identify that TDP-43 phase separation is regulated by acetylation, proteolysis, and ATPase-dependent chaperone activity of HSP70.

**One Sentence Summary:** Acetylation of TDP-43 drives its phase separation into spherical annuli that form a liquid-inside-a-liquid-inside-a-liquid.

---

A seminal discovery in the last decade has been recognition that large biological molecules, especially RNA binding proteins, can undergo <u>l</u>iquid-<u>l</u>iquid <u>p</u>hase <u>s</u>eparation (LLPS), resembling oil droplets in vinegar. Under certain physical conditions, proteins, nucleic acids, or a mixture of both in a complex solution can form two phases, a condensed de-mixed phase and more dilute aqueous phase (*1*). Inside the cell, proteins and/or nucleic acids de-mix into a condensed phase to form membraneless organelles which have been proposed to promote biological functions (*2*). One membraneless organelle is the nucleolus, first described in the early 1800’s and now recognized to be composed of proteins that undergo LLPS (*3–5*). Discovery that P-granules are de-mixed compartments with liquid behaviour brought widespread recognition to LLPS and its ability to mediate subcellular compartmentalization in a biological context (*6*). Phase separation has been also proposed for heterochromatin and RNA bodies (*1, 6, 7*). LLPS potentially underlies the operational principle governing formation of important organelles and structures, such as centrosomes, nuclear pore complexes, and super enhancers (*8–10*).

Few mechanisms to modulate intracellular LLPS have been identified. Disease-causing mutations of proteins such as FUS (*11*) and HNRNPA2B1 (*12*) alter their LLPS behaviors *in vitro,* but the physiological or pathological relevance of their intracellular LLPS has not been fully elucidated. Random, multivalent interactions among intrinsically disordered, <u>l</u>ow <u>c</u>omplexity <u>d</u>omains (LCDs) of proteins are a major driving force for LLPS *in vitro.* Oligomerization of other domains is thought to be required for LLPS *in vivo* (*13*). The inherent randomness of multivalent interactions (*14*) favors a model of disordered alignment – a conventional liquid phase in which molecules randomly move. The evidence is compelling that multiple material states can co-exist in the same de-mixed compartment, but how they assemble and whether there is order within demixed compartments is not determined. An exception is the liquid crystal, which flows like a liquid, but contains subdomains in which molecules are aligned (*15*).

Mislocalization and aggregation of the <u>h</u>eterogeneous <u>n</u>uclear <u>r</u>ibo<u>n</u>ucleo<u>p</u>rotein (hnRNP) <u>T</u>AR <u>D</u>NA-binding <u>p</u>rotein 43 (TDP-43) is a common pathological hallmark shared by several age-related neurodegenerative diseases, including amyotrophic lateral sclerosis (ALS) (*16*), frontotemporal dementia (FTD) (*15*), Alzheimer’s disease (AD) (*17*), and the recently defined AD-like variant named “LATE” (<u>l</u>imbic predominant <u>a</u>ge-related <u>T</u>DP-43 <u>e</u>ncephalopathy) in the oldest individuals (*18*). TDP-43 functions in pre-mRNA maturation, including splicing (*19*). However, in disease, heavily phosphorylated and ubiquitylated aggregates of TDP-43 accumulate cytoplasmically (*20*), accompanied by its nuclear clearance. We recently reported that TDP-43 naturally de-mixes in the nucleus, and shuttles to the cytoplasm in response to multiple stresses (*21*). Loss of nuclear TDP-43 causes usage of cryptic splice (*22–24*) and polyadenylation sites (*24*), dysregulating many important neuronal genes such as HDAC6 (*25*) and stathmin-2 (*22, 24*). RNA-binding by TDP-43 is regulated post-translationally (*26*) by acetylation of two lysine residues (K145 and K192) which inhibits its RNA-binding (*27*). Additionally, acetylated TDP-43 accumulates in cytoplasmic aggregates in the nervous systems of ALS and FTD patients (*27*).

Here we report a new LLPS phenomenon in which post-translational acetylation drives RNA-binding inhibited TDP-43 into <u>i</u>ntranuclear <u>L</u>iquid <u>S</u>pherical <u>A</u>nnuli (iLSA), forming a “liquid-inside-a-liquid-inside-a-liquid”, with the annuli comprised of a close-to-perfect liquid spherical shell with a higher TDP-43 concentration and an inner liquid spherical core with a lower TDP-43 concentration but enriched in HSP70 family chaperones.

## Results

### RNA binding-deficient TDP-43 de-mixes into iLSA in physiological conditions

Recognizing earlier work demonstrating that endogenous TDP-43 naturally phase separates in several cell culture models and in the mouse nervous system (*21*) and that an RNA-binding deficient TDP-43 promotes phase separation and aggregation in multiple cellular models (*27–29*), we tested if interaction between RNA and TDP-43 regulates LLPS of intranuclear TDP-43. TDP-43 contains an N-terminal self-association domain (*30, 31*), a C-terminal LCD, and two conserved <u>R</u>NA <u>r</u>ecognition <u>m</u>otifs (RRM1 and RRM2) whose RNA-binding activity can be abolished by acetylation of two lysine residues (K145 in RRM1 and K192 in RRM2) (*27*). We initially established that fluorescently tagged TDP-43 normally undergoes cycles of acetylation/deacetylation when expressed in non-neuronal (Fig. 1A) or neuronal (Fig. 1B) cell lines, as addition of the pan deacetylase inhibitors trichostatin A (TSA) or vorinostat (a.k.a. SAHA) was sufficient to drive TDP-43 into intranuclear, spherical annular droplets in which TDP-43 was enriched in the annular shell surrounding an apparently “hollow center”. Both the numbers and diameters of these annuli were enhanced by transient, partial inhibition of proteasome activity by addition of the FDA-approved bortezomib (BTZ) (Fig. 1B). Similarly, increasing acetylation by expression of a known acetyltransferase [the core component of CEBP-binding protein (CBP) (*32, 33*)] was sufficient to drive fluorescently tagged TDP-43 into spherical droplets or annuli, whose number increased following SAHA-mediated inhibition of deacetylases (Fig. S1A).

**Fig. 1.**
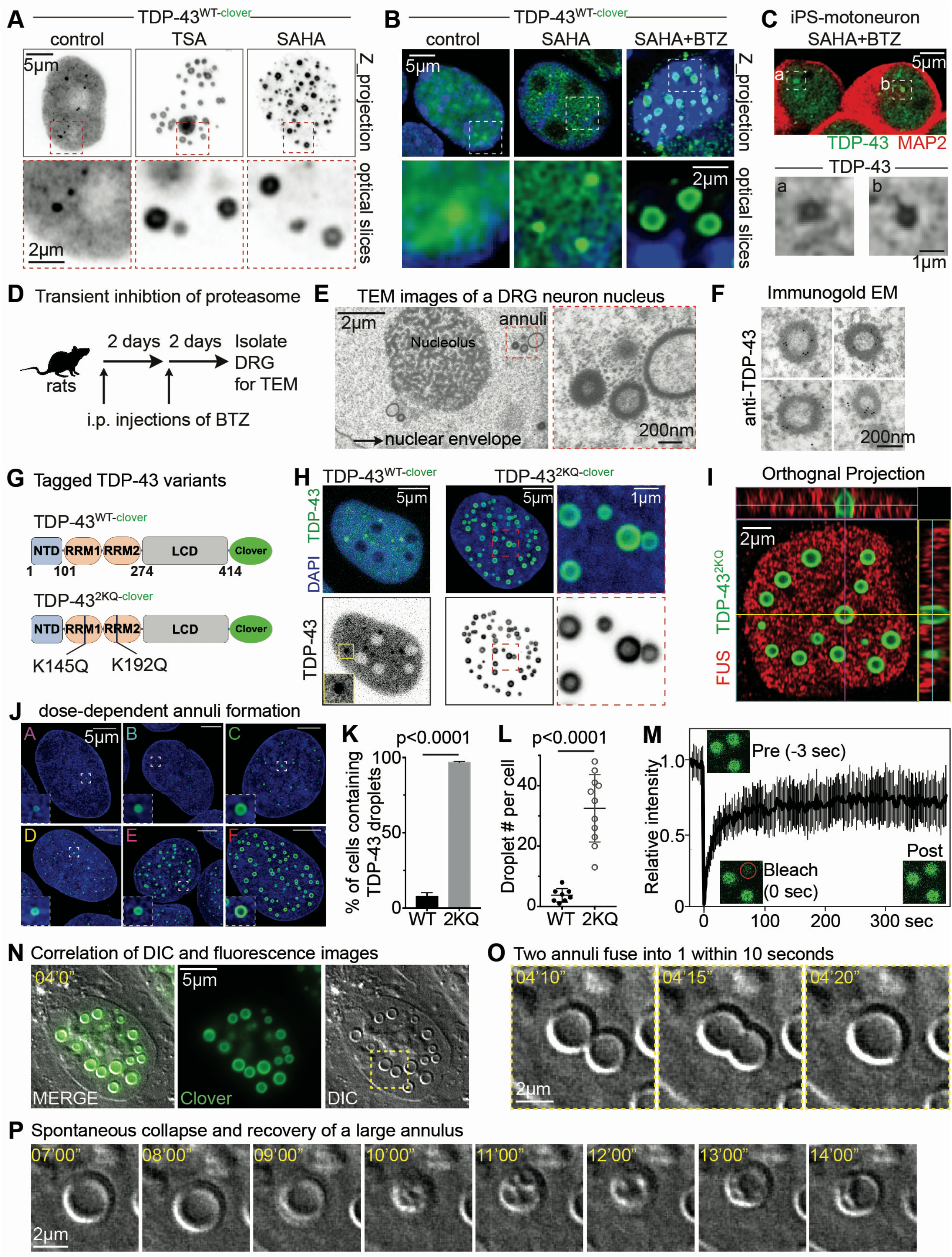
RNA-binding deficient TDP-43 naturally de-mixes into liquid spherical annuli which contains a liquid spherical core. (**A**) HDAC inhibitors TSA or SAHA induce nuclear TDP-43^clover^ to form annular droplets in HEK293T cells. Synergistic inhibition of HDAC and proteasome produce annular TDP-43 droplets in neuron-like SHSY-5Y cells (**B**), and in iPSC-derived motor neurons (**C**). (**D**) A scheme of inducing proteolytic stress in Sprague Dawley rats by intravenous (i.v.) injection of bortezomib saline. (**E**) Intranuclear spherical annuli were found in DRG sensory neurons under proteolytic stress. The red square is magnified. (**F**) Bortezomib-induced annuli contains TDP-43. (**G**) A diagram of wildtype and acetyl-mimic variant of TDP-43 (K145Q/K192Q, marked as 2KQ hereafter). Both proteins were tagged with a bright GFP variant, clover at the carboxyl-terminus. (**H**) U2OS cells expressing TDP-43^2KQ-clover^ exhibited much more nuclear granules than cells expressing TDP-43^WT-clover^. Upper panels: An example of U2OS cells inducible expression of clover-tagged TDP-43^WT^ and TDP-43^2KQ^. DAPI was used to stain nuclear DNA. The lower panels use monochrome to illustrate the compartmentalization of TDP-43. (**I**) Orthogonal projection demonstrates that TDP-43^2KQ-clover^ droplets are spherical annuli. (**J**) Annulus formation is dose dependent. Expression level of each clone is determined in Fig S1. (**K**) A quantitative analysis of the percentage of cells containing obvious TDP-43 droplets in U2OS cells expressing TDP-43^WT-clover^ or TDP-43^2KQ-clover^. (**L**) Number of nuclear droplets in U2OS cells expressing TDP-43^WT-clover^ or TDP-43^2KQ-clover^. (**M**) Intranuclear TDP-43^2KQ-clover^ spherical annuli are liquid-like compartments, demonstrated by <u>f</u>luorescence <u>r</u>ecovery <u>a</u>fter <u>p</u>hotobleaching (FRAP) experiments. (**N**) The dynamic fusion of TDP-43^2KQ-clover^ spherical annuli was directly visualized by Differential Interference Contrast (DIC) microscopy. Two annuli (1~2μm) fused into 1 annulus (>2 μm) in 10 seconds, observed by live imaging (**O**). (**P**) A series of images shows the collapse and recovery of a big annulus (>2 μm). The big annulus collapsed into a droplet containing multiple droplets, which then fused into a new annulus in 5 minutes.

Annulus formation after deacetylase inhibition coupled with transient proteasome inhibition did not require elevated TDP-43 accumulation: wild type TDP-43 expressed from the endogenous alleles assembled into similar spherical intranuclear annuli in induced pluripotent stem cell (iPSC)-derived human motor neurons (Fig. 1C). Perhaps most remarkably, partial proteasome inhibition *in vivo* in rat dorsal root ganglion neurons (after peripheral injection of BTZ) produced easily observable spherical annuli (0.2 – 1 μm in diameter) whose annular shells were enriched in endogenous, wild type TDP-43 (as determined using immunogold electron microscopy) (Fig. 1D-F).

To determine how TDP-43 acetylation affected its LLPS, we next used an established (*27*) TDP-43 acetylation-mimic TDP-43^K145Q,K192Q^ (to be referred to hereafter as TDP-43^2KQ^) in which glutamines replace the two lysines that are acetylated in TDP-43 (Fig. 1G). After stable lentiviral-mediated integration of doxycycline-inducible TDP-43 genes (Fig. S1B,C), clones were isolated that upon induction accumulated fluorescently tagged TDP-43^2KQ^ between 0.3 and 9 times endogenous TDP-43 (Fig. S1D-G). RNA-binding deficient TDP-43^2KQ^ formed close-to-perfectly round annuli in which it was enriched in the annular shells and diminished in the centers, as revealed by confocal microscopy (Fig. 1H) or orthogonal projection of high-resolution confocal image stacks (Fig. 1I). Annuli numbers and diameters increased in a TDP-43 dose-dependent manner (Fig. 1J). At high levels of accumulation, over 95% of cells expressing TDP-43^2KQ^ contained 20-50 large (0.5~2 μm in size) annuli (Fig. 1K), compared with <10% of cells expressing a comparable level of fluorescently-tagged TDP-43^WT^. Cells expressing TDP-43^2KQ^ contained >6 times more annuli than those with TDP-43^WT^ (Fig. 1L).

<u>F</u>luorescence <u>R</u>ecovery <u>A</u>fter <u>P</u>hotobleaching (FRAP) was used to determine that most RNA binding-deficient TDP-43^2KQ^ in annuli displayed liquid-like behavior, freely exchanging with the nucleoplasm with a recovery half time of 9 sec. (Fig. 1M). Use of 3D live cell imaging of intranuclear annuli (unambiguously identified as annuli by Z projections - Fig. S1H and Movie S1) containing fluorescently tagged TDP-43^2KQ^ confirmed that the annuli frequently fused to form larger annuli (Fig. S1I). Furthermore, direct comparison of fluorescence and differential interference contrast (DIC) imaging revealed that annuli could be directly observed in DIC images (Fig. 1N). Live cell DIC imaging (Movie S2) detected abundant, rapid (within 10 seconds) annuli fusion events in which the shells fused with each other, as did the cores, behavior indicating that both shells and cores behaved like liquids (Fig. 1O and Movie S2). Annulus fission was not observed; instead, the largest annuli collapsed into irregular, multi-compartmental droplets that with time recovered into annuli again (Fig. 1P). Thus, both the shell and core of annuli formed by RNA binding-deficient TDP-43 display liquid properties. We will refer to these as intranuclear <u>l</u>iquid <u>s</u>pherical <u>a</u>nnuli (iLSA).

### iLSA from RNA binding deficient TDP-43 recruit wild type TDP-43

We tested how iLSA containing RNA binding deficient TDP-43 affected behavior of wild type TDP-43. We started with a cell line (*21*) in which lentiviral transduction had been used to functionally replace endogenously expressed TDP-43 with a fluorescent, mRuby2-tagged, wild type TDP-43 (hereafter TDP-43^WT-mRuby^), with TDP-43 autoregulation (*19, 34*) maintaining overall TDP-43 level (Fig. S2A-C). We then introduced a previously characterized RNA binding deficient TDP-43 variant (*35*) in which five phenylalanine residues were mutated in the two RRM domains to produce TDP-43^5FL-clover^. As expected, TDP-43^5FL-clover^ de-mixed into intranuclear spherical annuli (Fig. S2D), behavior indistinguishable from that previously documented for TDP-43^2KQ^. Over an 18 hour time course, TDP-43^WT-mRuby^ was recruited into annuli indistinguishably from TDP-43^5FL-clover^ (Fig. S2E,F). Use of FRAP to bleach either the RNA binding mutant TDP-43^5FL-clover^ or TDP-43^WT-mRuby^ within half of a nucleus revealed comparably rapid kinetics of recovery of red and green fluorescence within both shells and cores of individual bleached annuli, with corresponding fluorescence losses from unbleached annuli (Fig. S2G,H). Thus, RNA binding deficient TDP-43 can effectively recruit wild type TDP-43 into the liquid shells and cores of intranuclear annuli.

### TDP-43 iLSA is a selective barrier excluding intranuclear components

Examination of iLSA from acetylation mimicking TDP-43 revealed the exclusion of several intranuclear components, including chromatin (marked by histone H2B - Fig. 2A), behavior similar to that previously reported for LLPS of artificial nuclear proteins and consistent with exclusion mediated by surface tension (*13*). Also excluded were some RNA-binding proteins, including FUS (mutations in which cause ALS) and iavNP, a virally-encoded RNA-binding protein (Fig. 2B). Other RNA-binding proteins (hnRNPA2B1, hnRNPH1 and hnRNPK) freely disused into both the shell and core of the annuli (Fig. 2C), while nuclear targeted GFP (*36*) was enriched in the core but depleted from the annular shell (Fig. 2B). Recognizing that iLSA contained both RNA binding deficient and wild type TDP-43, as well as other RNA binding proteins, we then determined annular RNA content by using CLICK-chemistry to fluorescently label RNA after incorporation of 5-ethynyl uridine (5-eU) (Fig. 2D). Imaging of individual annuli and surrounding nucleoplasm revealed that RNA was depleted from annular shells and cores relative to nucleoplasm (Fig. 2E,F). Remarkably, annuli remained intact after exposure to the cytoplasm during mitosis (Fig. 2G) and were then excluded from newly forming post-mitotic nuclei in earliest interphase (as determined by live cell imaging - Fig. 2H). Cytoplasmic TDP-43^2KQ^ was imported into newly formed nuclei and reassembled into iLSA within 2 hours after mitotic exit (Fig. 2H). Analysis of annuli in cells containing a micronucleus produced by a chromosome segregation error revealed a time-dependent post-mitotic increase in the number of annuli, presumably assembled spontaneously from TDP-43 reimported through nuclear pores (Fig. 2H).

**Fig. 2.**
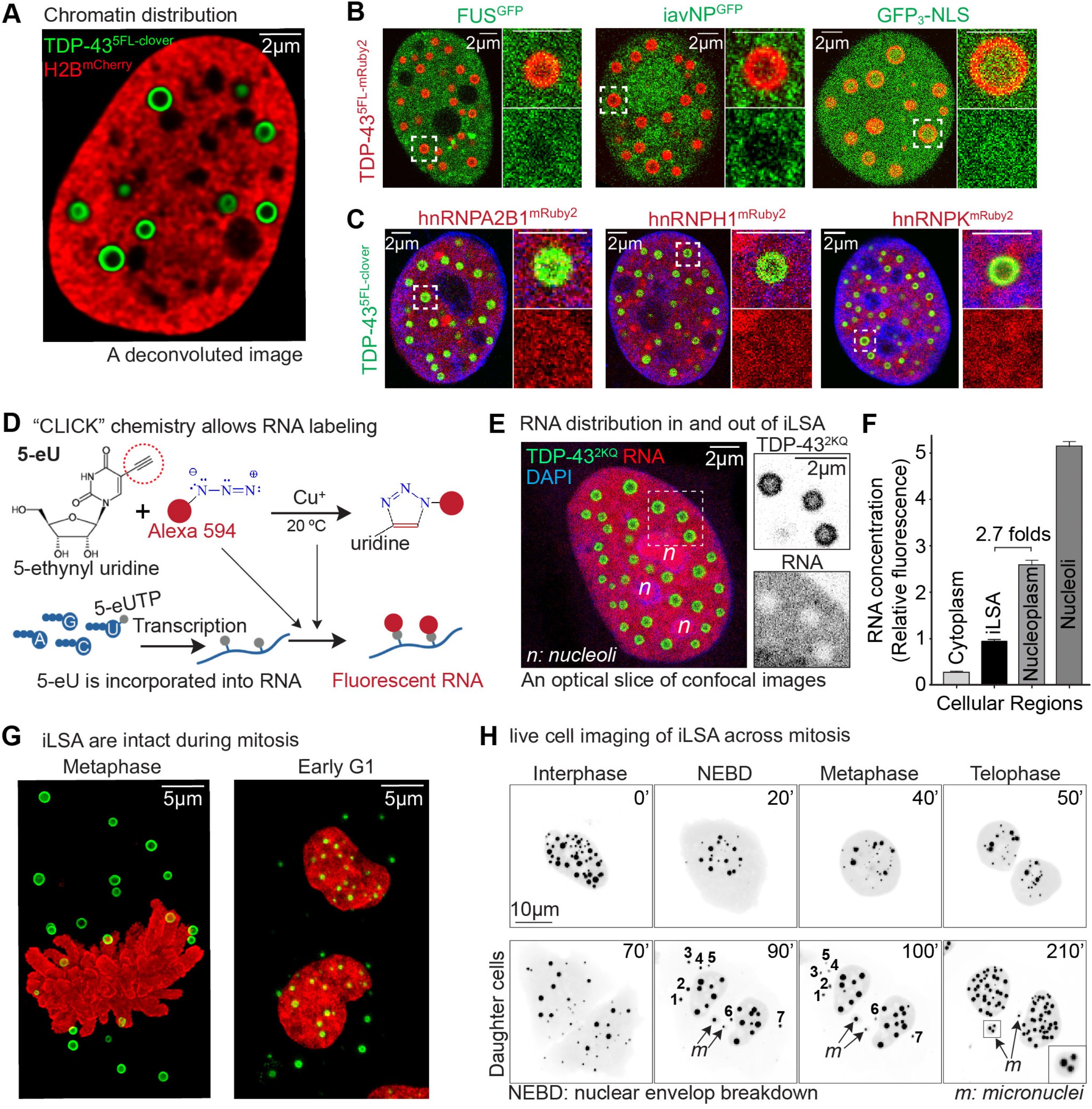
TDP-43 iLSA are selective barriers for nuclear proteins and RNA. (**A**) A U2OS nucleus stably expressing a fluorescent histone marker, H2B^mCherry^, and TDP-43^2KQ-clover^. (**B**) Nuclear RNA-binding proteins hnRNPA2B1, hnRNPH1 and hnRNPK show even distribution in and out of the annuli. Each protein is mRuby2 tagged at C-terminus. (**C**) EGFP-labeled FUS and iavNP are decreased in the annuli, while EGFP alone show even distribution in and out of the annuli. (**D**) Cellular RNA was fluorescently labeled by “CLICK” chemistry cells. (**E**) Nuclear RNA (labeled by CLICK chemistry) was largely excluded from the annuli. (**F**) Quantification of fluorescence intensity of the RNA. (**H**) An image series shows the dynamics of TDP-43^2KQ-clover^ annuli during mitosis.

### TDP-43 iLSA are densely packed nuclear compartments

Conventional transmission electron microscopy revealed that despite the liquid character of shells and cores, iLSA were easily visualized (with a standard OsO4 staining protocol) following induction of RNA binding deficient TDP-43^2KQ^, but not TDP-43^WT^ (Fig. 3A-C). Cross sections of TDP-43 annuli were close-to-perfect circles of diameters between 0.5 and 2 μm (Figs. 3C and S3A-F). Distribution of TDP-43^2KQ-clover^ molecules across the annuli was determined using 70~80nm ultrathin sections of fixed cells incubated with gold particle-labeled GFP antibody recognizing Clover. Antibody-bound gold particles were highly enriched in the annuli, slightly enriched in the annular core, and very sparse in the nucleoplasm (Figs. 3D and S3). In contrast, TDP-43^WT-clover^ formed droplets in which TDP-43 was uniformly distributed (Fig. 3D). Some annuli contained electron-lucent regions in the electron-dense annuli, consistent with fusion events or a liquid phase exchanging between the inner droplet and the nucleoplasm (Fig. S3G). Average annular shell thickness (Fig. S3H) was independent of the annulus diameter (Fig. S3I).

**Fig. 3.**
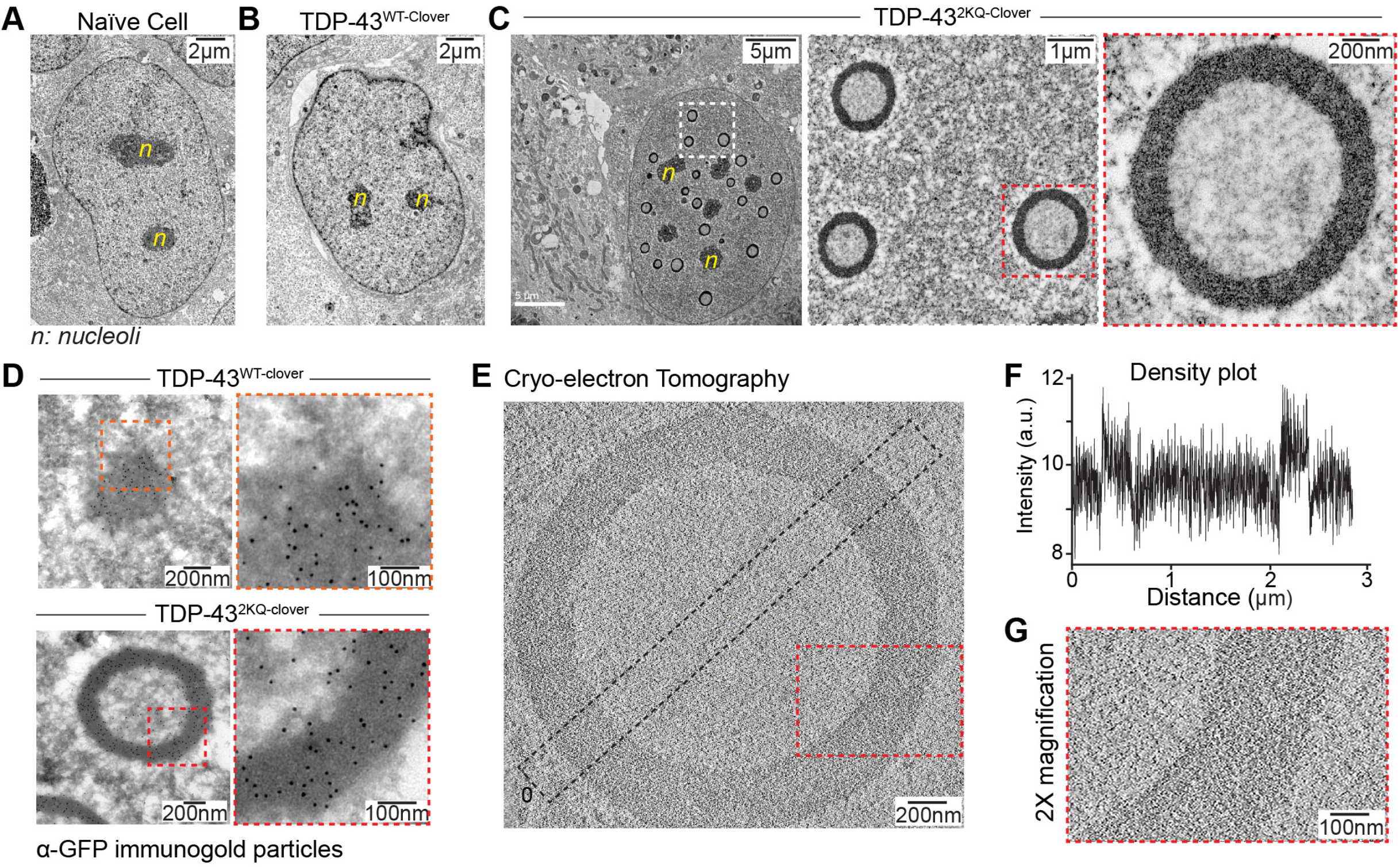
TDP-43 iLSA are densely aligned membraneless compartment that can be directly identified by Transmission electron microscopy (TEM) and cryo-electron tomography (CryoET). TEM examples of nuclei from a naïve U2OS cell(**A**), a U2OS cell expressing TDP-43^WT-clover^ (**B**), a U2OS cell expressing TDP-43^2KQ-clover^ (**C**). More examples are shown in Fig. S3. (**D**) Immunogold particles labels TDP-43^WT-clover^ droplet or an TDP-43^WT-clover^ annulus. The dashed boxes are enlarged to show the distribution of gold particles. Enlarged examples are shown in Fig. S4. (**E**) CryoET of an annulus. A density plot of the black box is shown in (**F**), and the red box is magnified in (**G**).

Cryo-electron tomography (CryoET) was used to determine annuli structure in physiological conditions (Fig. 3E). After cryo-FIB milling to create thin lamellae from cells (Fig. S5), tomographic reconstructions revealed that annuli had a well-defined annular ring of 320 nm ± 20 nm thickness, consisting of meshwork-like densities similar to the meshwork reported in tomographic images of *in vitro* reconstituted phase-separated compartments (*37*). Molecular density in the annular core appeared lower than in adjacent nucleoplasm. The tomograms reveal annular droplets consistent with a liquid center surrounded by a porous, meshwork-like rim through which molecules can shuttle.

### The nuclear TDP-43 spherical annulus exhibits liquid crystalline properties

Liquid crystals flow like a conventional liquid but contain non-randomly oriented molecules producing a birefringent, anisotropic liquid (*15*). To determine if the apparently densely packed molecules in iLSA are anisotropic, we used complete extinction microscopy with a pair of polarizers perpendicular to each other (Fig. 4A,B), conditions in which image brightness correlates with the level of anisotropy (*38*). Annular shells (but not cores) were visible, indicating that some orderly aligned molecules form anisotropic subdomains in the internal structure (Fig. 4C, Movie S3). Anisotropic lipid bilayers of all cellular membrane compartments, another biological liquid crystal (*39*), were also visible under the complete extinction condition (Fig. 4C, the left panel), while in contrast, naturally phase separated nucleoli (*5*) imaged with extinction microscopy were not distinguished from the surrounding nucleoplasm (Fig. 4C-D). In fact, iLSA were the only visible structures formed by proteins inside cells under complete extinction microscopy. Live imaging (Fig. 4E) provided further evidence of the liquid properties of the shells, with the positions of the ordered subdomains changing faster than the (180 ms) detection limit.

**Fig. 4.**
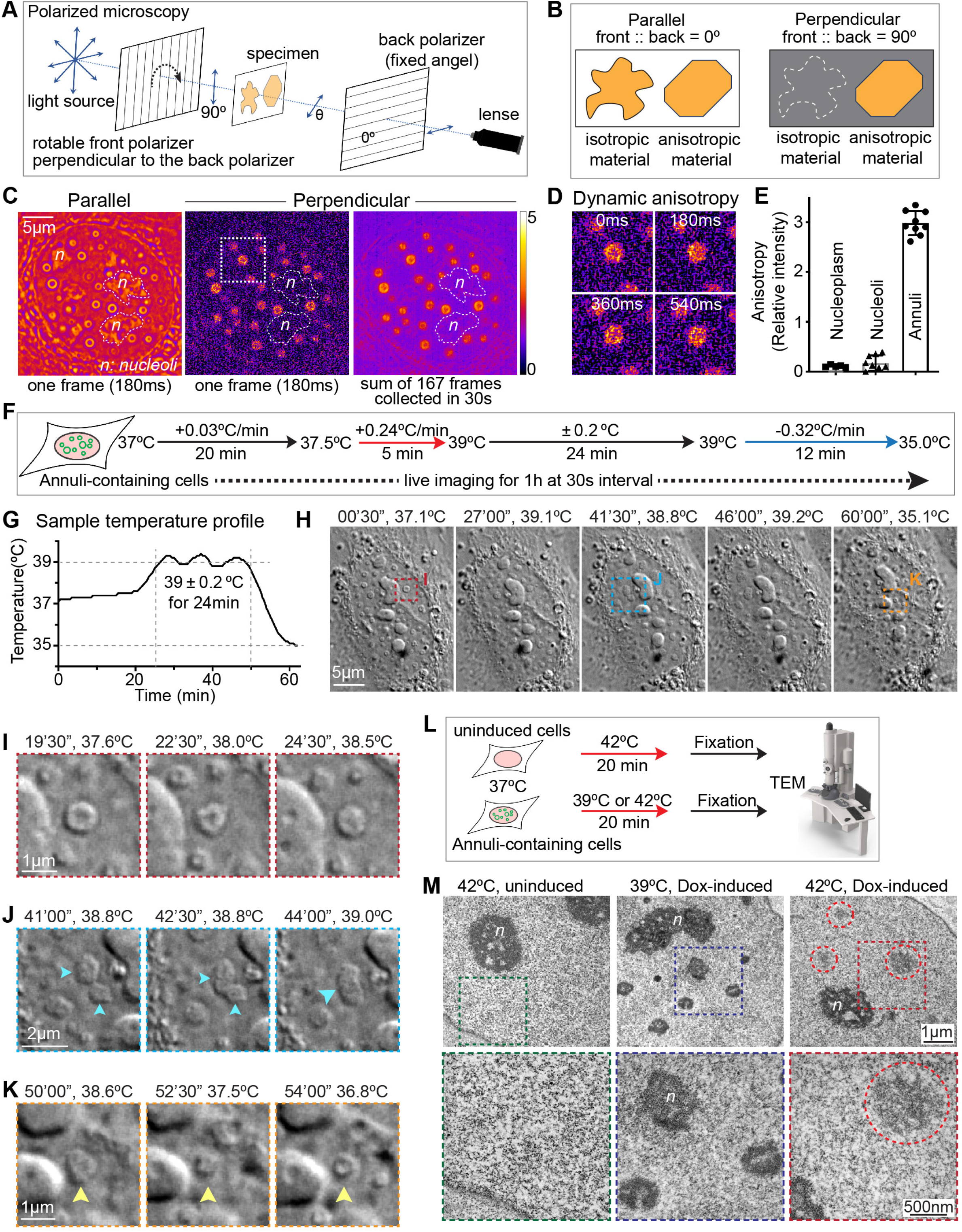
TDP-43 iLSA exhibit liquid crystalline properties. (**A**) A schematic explanation of complete extinction microscopy. The light path contains two polarizers: the front polarizer can be adjusted from 0° to 90°, and the back polarizer has a fixed angle. Rotating the front polarizer result in a change of polarized light. (**B**) Isotropic and anisotropic materials can be distinguished by complete extinction microscopy because the birefringent property of the anisotropic material allows some light to pass the back polarizer. (**C**) The same nucleus containing TDP-43 annuli under parallel (the left panel) or perpendicular (the middle panel) conditions. The right panel demonstrated by a projection of 167 frames collected in 30 seconds. The phase-separated nucleoli (marked by “*n*”) cannot be detected under the complete extinction microscopy. Gray scale images are pseudo-colored to show better contrast. (**D**) Dynamic anisotropic sub-domains were observed by live imaging. This image series shows an enlarged annulus highlighted in the white square in panel **c**. (**E**) Quantification of the intensity of nucleoplasm, nucleoli and spherical annuli in three cells expressing TDP-43^2KQ-clover^. (**F**) Experimental design of live cell imaging performed under a continuous temperature shift. (**G**) The sample temperature profile measured during the temperature shift. (**H**) A DIC image series shows an annuli-containing nucleus under different temperature during the time shift. (**I-K**) Enlarged imaging series at different time point highlighted in panel **H**. (**I**) shows an annulus (highlighted in the red square) become uniform droplet as temperature increased from 37.6°C to 38.5°C. (**J**) shows the fusion of two uniform droplets (highlighted in the blue square) at 39°C. (**K**) shows a uniform droplet became a spherical annulus as the temperature cooled from 38.6°C to 36.8°C. (**L**) A schematic explanation of 20-minute temperature shift and determination of the structural changes by TEM. (**M**) The temperature shift-induced structural change was confirmed by TEM. Irregular electronic dense compartments were circled in red.

Anisotropy of ordered biomacromolecules changes or disappears as temperature increases (*40, 41*) because increased entropy changes the interaction between the biomacromolecules that mediate the anisotropic alignment. Consequently, we performed live DIC imaging during temperature increase from 37 to 39 °C (Fig. 4F-H, Movie S4). Annuli gradually transformed into uniform liquid droplets (Fig. 4I and S7B) which fused and changed shape over time (Fig. 4J and S6C). Uniform droplets reconverted into annuli as the temperature was reduced back to 37°C (Fig. 4K and S6D). Temperature-dependent conversion from annuli to liquid droplets was confirmed by electron microscopy. Extended time at elevated temperature (20 minutes at 39°C) partially disrupted annular structure, producing more irregular shapes, while after shift to 42°C the shell disappeared, yielding uniform droplets of lighter electron density (Fig. 4M).

### Oligomerization and prion-like domains of TDP-43 are required for annular phase separation

To identify the interactions that mediate annulus formation, we first determined that forcing RNA binding deficient TDP-43 accumulation in the cytoplasm (by mutation of the nuclear localization sequence) eliminated iLSA even in the most highly expressing cells (Fig. S7B). TDP-43 possesses an N-terminal self-interaction domain (NTD) (*30*) that mediates its oligomerization (Fig. S7C,D) through three intermolecular salt bridges (*30, 31*). Disrupting self-interaction [by converting serine residue 48 in one bridge to a phosphorylation mimicking residue (Fig. S7E,F) or by mutating glycine residue 53 that is part of forming a second bridge (Fig. S7E,F)] completely disrupted annulus formation. The NTD alone did not form intranuclear droplets or annuli, but was sufficient to drive intranuclear phase separation into droplets, but not annuli, when linked to LCD domains from FUS or EWS (Fig. S7G,H), other RNA binding proteins previously shown to be able to phase separate (*11*).

### Modeling identifies that a self-associating TDP-43 partner can drive annulus formation

To identify mechanisms that can drive annulus formation, we created a mathematical model of an RNA binding deficient TDP-43 in the presence of RNA (as it would be in a cell). Because the majority of TDP-43 molecules form stable dimers or even oligomers *in vitro (31, 42)* and perhaps *in vivo (30, 43)* and because nuclei have a high RNA concentration (Figs. 2F, 5A, and S8), we modeled a de-mixing system which includes RNA and acetylated TDP-43 monomers and dimers, neither of which have affinity for the RNA. We employed a Cahn Hilliard diffuse interface model coupled with a Flory Huggins free energy scheme that employs the parameter χ to describe the energetic favorability of different components mixing in the system. Each pair of solute species has a parameter χ representing their energy cost of mixing. Higher values of χ represent a larger energy cost, favoring de-mixing. In this model, two acetylated TDP-43 monomers (P) come together to form the TDP-43 dimer (P_2_). We further assumed that once formed, it was energetically unfavorable for the dimer (P_2_) to mix with both the nucleoplasm and the RNA. Conversely, RNA was diffusely distributed throughout nucleoplasm. With this modeling structure, acetylated TDP-43 monomers formed dimers (P_2_) which then rapidly de-mixed from the RNA-containing nucleoplasm forming only uniform droplets, not annuli (Fig. S8D).

**Fig. 5.**
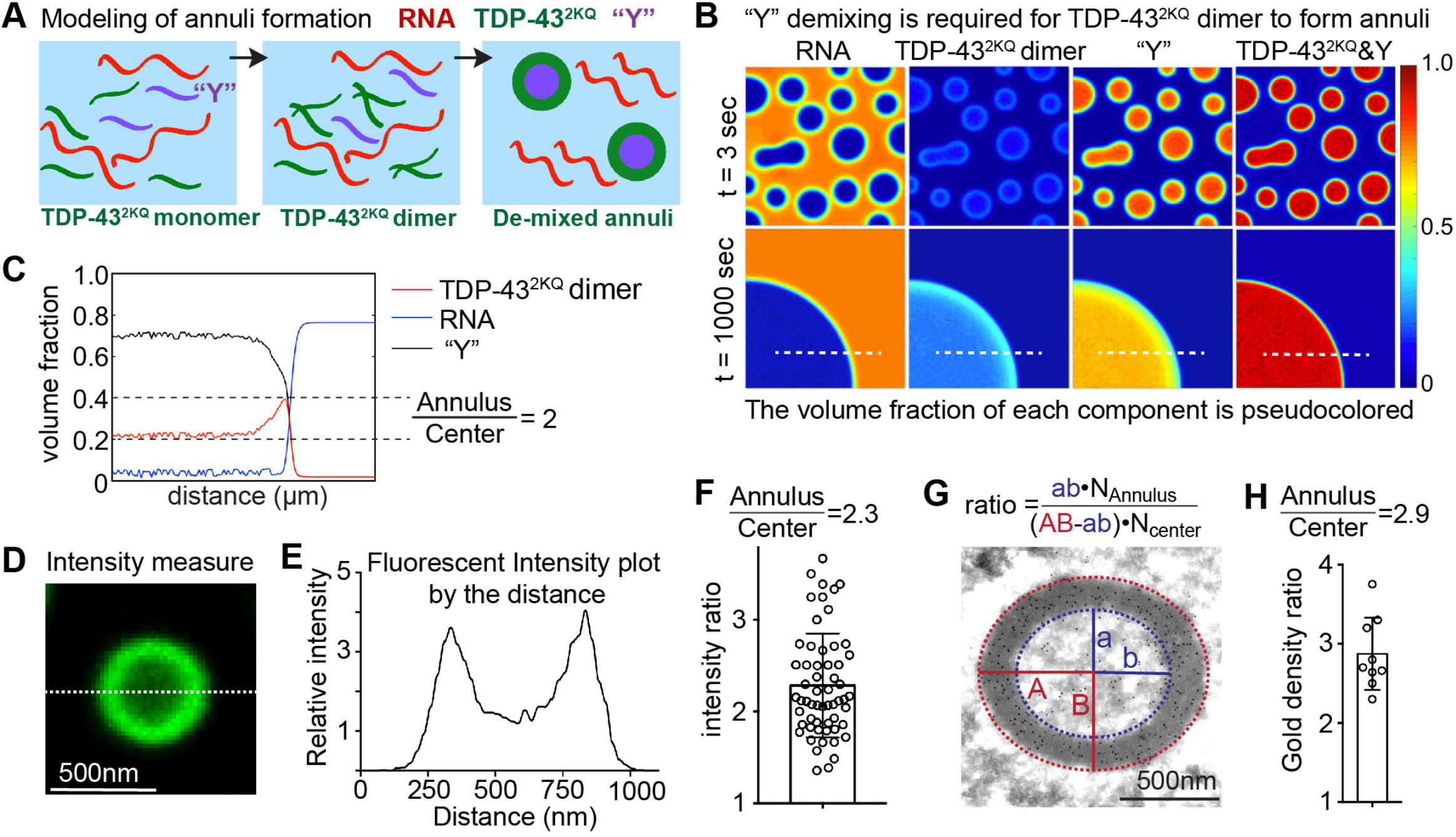
Mathematical modeling predicts the liquid core components are required for annulus formation. (**A**) A scheme explains the modeling of annulus formation by TDP-43^2KQ^ mutant in a simple mixture composed of TDP-43, RNA and Y molecules. (**B**) Snapshot of modeled de-mixing at 3 and 1000 second after the starting time point. (**C**) The intensity measurement of modeled de-mixing at 1000 second time point. The ratio of volume fraction between the annuli and the center droplet for TDP-43^2KQ^ is about 2. (**D, E**) An example of fluorescence intensity measurement from a raw data of optical slices. (**F**) Quantification of fluorescence intensity ratio between the annulus and the center. The mean is 2.28, and the standard deviation is shown. (**G**) Measuring the ratio of TDP-43^2KQ-clover^ distribution by the gold particle density in the immunogold-labeling experiments. (**i**) Quantification of the ratio of gold particle density between the annulus and the center.

Because a simple mixture of acetylated TDP-43 and RNA failed to form annuli in our initial model, we extended it to include a hypothetical entity, Y, representing one or more proteins that possess both self-interaction and propensity to de-mix with TDP-43 dimer. This approach built upon prior modeling work in which two components from a surrounding solution de-mixed into the same droplet, with one occupying the spherical core and the other forming an annulus between the droplet core and the diffuse phase (*44*). Upon introducing this hypothetical Y protein into the model, annular de-mixing was induced, but only when TDP-43 dimer bound Y with an intermediate affinity (Figs. 5B and S9). Further, because RNA is not enriched inside the TDP-43 annuli (Fig. 2D), we also hypothesized that it was energetically unfavorable for Y to mix with the diffuse RNA. Introduction of Y into a mixture of acetylated TDP-43 and RNA successfully modeled annular de-mixing of TDP-43, with Y forming the annular core (Fig. 5B), with the predicted volume fraction of TDP-43 in the annular shell about two times that of the inner droplet (Fig. 5C). The model closely predicted the actual 2.3-2.9 fold experimentally measured ratio of TDP-43 in shell and core of annuli formed with TDP-43^2KQ-clover^ [determined by fluorescence intensity from optical slices (Fig. 5D-F) or by immunogold electron microscopy (Fig. 5H-I)].

### HSP70 chaperones drive iLSA by forming the inner liquid core with TDP-43

Because our introduction of “Y” allowed iLSA formation in our model, a proteomic approach was undertaken to determine the identity(ies) of such components in the liquid cores. A combination of differential isotope labeling of cells with or without TDP-43 iLSA was coupled with proximity labeling (with biotin) of proteins close to peroxidase tagged TDP-43 (Fig. S10A) followed by immunoprecipitation and quantitative mass spectrometry (Fig. 6A). When stably expressed, TDP-43^mRuby2-APEX^, a wildtype TDP-43 fused to red fluorescent protein (mRuby2) and peroxidase (APEX2) tags, accumulated intranuclearly with a proportion de-mixed into droplets (Fig. S10B). Induction of Clover-tagged RNA binding deficient TDP-43^2KQ^ yielded co-de-mixing of the two TDP-43’s into abundant intranuclear annuli (Fig. S10B). Cells were isotope-labeled in the absence or presence of annuli and briefly exposed to H_2_O_2_ to enable APEX2 to label proximal proteins with biotin. After immunopurification of biotinylated proteins, quantitative mass spectrometry (Fig. 6B) was used to identify that only 5 proteins were enriched at least 6 fold in TDP-43-containing annuli. All 5 (HSPA1A, HSPA1L, HSPA5, HSPA6, and HSPA8), with enrichments in annuli of 6-16 fold, are members of the HSP70 family of ATP-dependent protein folding chaperones (Fig. 6B). Since we determined that indirect immunofluorescence cannot be used to determine content of the cores because conventional fixation produces a barrier of crosslinked annular shell proteins that prevent antibody penetration to the core (Fig. S11), we used expression of mRuby2 tagged HSPA1A, HSPA1L, HSPA6 and HSPA8 to validate that each HSP70 family member was indeed enriched (by 3-10 fold) in the cores of annuli formed with RNA binding deficient TDP-43^2KQ^ (Fig. 6C).

**Fig. 6.**
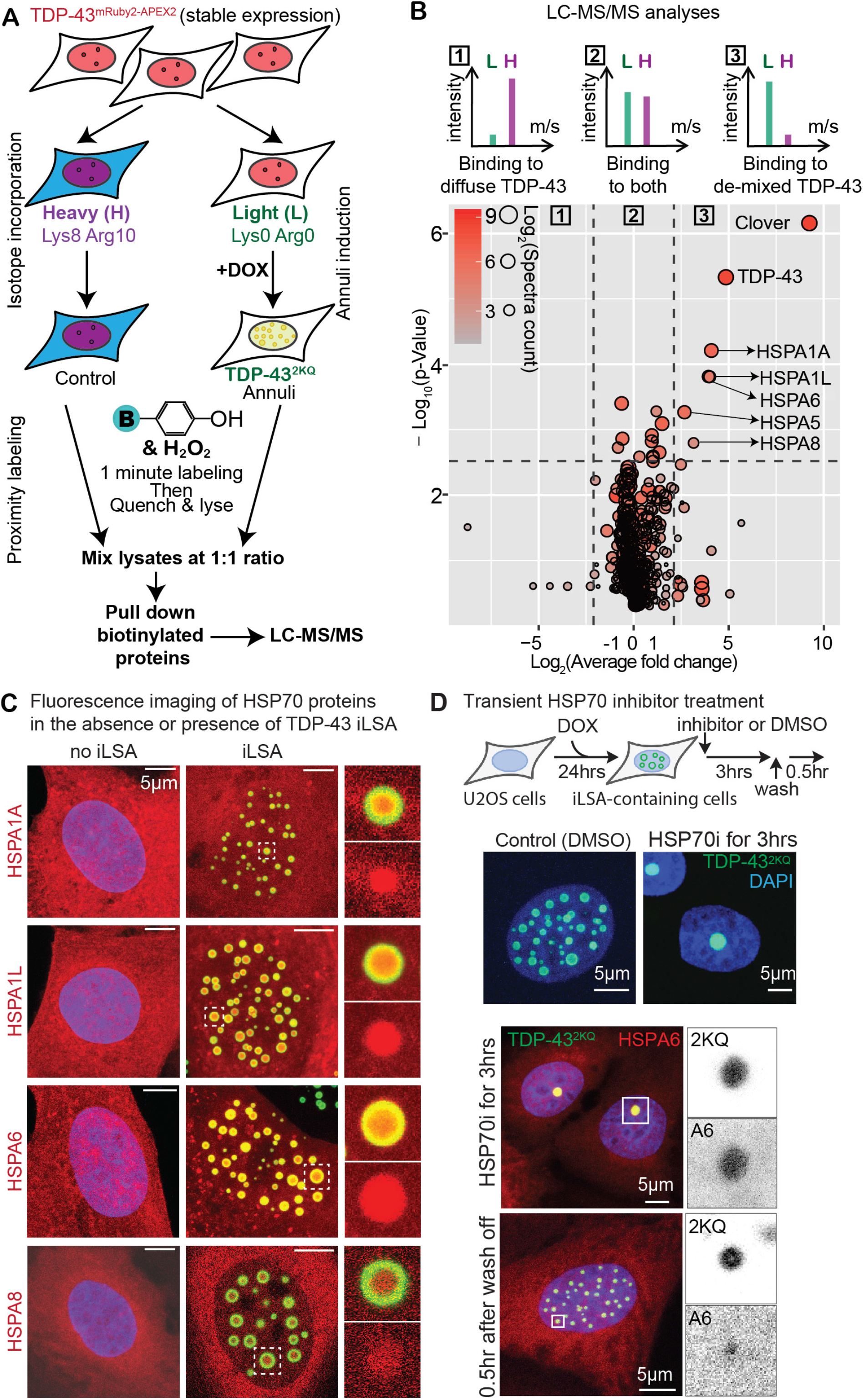
The HSP70 family Chaperones are the major components enriched in the liquid core. (**A**) A schematic explanation of APEX2-mediated proximity labeling followed by quantitative proteomic analyses. The mechanism of proximity labeling is shown in Fig. S10A. (**B**) Quantitative proteomic analyses identify HSP70 family proteins enriched in the de-mixed TDP-43. Y axis is p-value at log_10_ scale, and X axis is fold enrichment at log_2_ scale. The size and color of each spot correspond to the total spectra count of LC/MS-MS reads at log_2_ scale. (**C**) Maximum intensity projections of immunofluorescence images of U2OS nuclei expressing TDP-32 annuli (green) and HSP70 family proteins tagged by red fluorescent protein mRuby2. Magnified Confocal optical slices of the white boxes are shown on the right side. (**D**) TDP-43 iLSA expressing cells form nuclear gels after inhibiting HSP70, which is reversible after washing off of the HSP70 inhibitor. Upper panels: compare to DMSO, 3-hour treatment of HSP70 ATPase inhibitor (VER155008) causes iLSA to form a big intranuclear gel. Lower panels: HS70i-induced nuclear gel contains TDP-43^2KQ^ and HSP70 uniformly distributed inside the nuclear gel, which become iLSA 30 minutes after removing the inhibitor.

Each HSP70 family member requires ATP hydrolysis to fulfill its protein folding chaperone function (*45*). After addition of an HSP70 inhibitor thought to block ATP hydrolysis by all family members (*46*) and to lock chaperone binding to a client protein (*45*), live imaging revealed that inhibition of HSP70 chaperone activity induced TDP-43^2KQ^-containing annuli conversion (within 10 minutes) into small droplets of uniformly distributed TDP-43^2KQ^ and HSP70, which then fused within an additional 30~60 minutes into a single large granule (Figs. 6D, S10D,E). These large nuclear granules were reversible, with iLSA reforming within 6 minutes after inhibitor removal (Fig. S10E).

## Discussion

Here we report that acetylation of TDP-43 induces it to phase separate into round, intranuclear annuli in which acetylated and unmodified TDP-43 are enriched in an anisotropic, liquid annular shell and the HSP70 family chaperones (as well as TDP-43) are the major components of a spherical liquid core, whose liquidity requires continuing ATPase-dependent chaperone activity. TDP-43 iLSA differ from previously identified intracellular liquid droplets, including the nucleolus (*47*), in that the two phases are well aligned to produce symmetrical annuli. TDP-43 iLSA exclude chromatin and some RNA binding proteins. Recognizing that protein acetyltransferases are enriched in transcription initiation complexes (*32*), phase separation of TDP-43 near sites of transcription could form a barrier to exclude some RNA-binding proteins but not others and allowing regulated RNA splicing. TDP-43 iLSA exhibit birefringence, evidence that an anisotropic liquid can be formed intracellularly by protein phase separation. While acetylation of histones H1 and H3 has recently been shown to abolish liquid-liquid phase separation of chromatin (*48*), acetylation of TDP-43 does the opposite: it promotes phase separation.

We have established the principles of annuli formation using mathematical modeling and experimental validation of the model’s predictions that phase separation into annuli is driven by TDP-43 self-association and a weaker affinity for another factor which we identify to be the HSP70 family of chaperones. As predicted, annular de-mixing is shown to produce 2-3 fold TDP-43 enrichment in the anisotropic liquid shells and a corresponding 5-16 fold enrichment of the HSP70 family molecular chaperones in the isotropic liquid cores. In our model, iLSA formation required a low affinity “Y” component, with annuli converting to uniformly phase separated droplets when Y was removed or its affinity to TDP-43 was increased. We experimentally validated the latter of these predictions, as we found (Fig. 4F-H) that the annular shell rapidly merged with the liquid core when the HSP70 affinity for TDP-43 was elevated by mild temperature increase (*49*) or by transiently locking the chaperones onto their clients by inhibition of their ATPase cycles (Fig. 6D) (*45*).

Our modeling strongly supports an essential role for the HSP70 chaperones in annuli formation and we have demonstrated that these chaperones play active roles in maintaining the liquidity of the internal annular core, as those centers reversibly convert into gels or solids when the HSP70 family ATPases are inhibited (Figs. 6D and S10D,E). Acetylation-mediated loss of RNA-binding could expose one or more unstable interfaces within TDP-43’s RRM domains that contain many hydrophobic residues that mediate protein-RNA interaction (*50*), which in turn recruit and are stabilized by HSP70 chaperone activity. While HSP70’s cytoplasmic activity has been reported to be required to disassemble stress granules (*51*), the combination of our modeling and experimental evidence shows that intranuclear HSP70 activity facilitates iLSA formation. We note that TDP-43 is the only hnRNP that contains a folded, self-interacting N-terminal domain, which is likely to enable its phase separation into anisotropic annuli. We further validated that the self-interaction of the N-terminal domain is mediated by salt bridges formed by residues Gly53 and Ser48 of one unit with Glu17 of the neighboring unit (*30*). N-terminal self-interaction can be abolished by phosphorylation on Serine 48 (Fig S7E) (*31*). However, the kinase has not been identified yet.

Annuli similar to those formed by acetylated TDP-43 have been reported in Drosophila germ granules, first reported as intranuclear spherical annuli (*52*) and later shown to be gel-like (*53*). Furthermore, annular nuclear bodies, apparently containing both polyA-RNA and the RNA binding protein SAM68, have also been reported (*54*) in degenerating motor neurons of SMNΔ7 mice, a model of spinal muscular atrophy type II. Matrin3, another ALS-causing gene, forms round annulus-like nuclear droplets when its second RNA-binding domain is removed, (*55*), albeit the liquid properties of these intranuclear droplets have not been determined. Here we show that nuclear TDP-43-containing annuli can be driven by altering the balance of ongoing acetylation/deacetylation (Fig. 1A-C) or in neurons of the mammalian nervous system that experience proteomic stress from transient proteasome inhibition (Fig. 1D-G).

Abnormal aggregation of TDP-43 is nearly a universal feature in ALS and in FTD with ubiquitin positive inclusions (FTLD-U) (*56, 57*). There is increasing evidence of TDP-43 pathology in the hippocampal neurons of patients with Alzheimer’s disease and correlated with ApoE4 alleles (*58–60*). LATE, a recently defined dementia in the oldest elderly, is a neurological disorder of remarkable TDP-43 cytoplasmic aggregates (*18*). Acetylated, RNA-binding deficient TDP-43 is present in the pathological TDP-43 aggregates (*27*). Moreover, intranuclear TDP-43 granules were observed in postmortem FTD patients (*61*). Because our evidence has shown that TDP-43 iLSA are sensitive to the ATP-dependent HSP70 activity, these intranuclear inclusions observed in patient may be formed from TDP-43-containing iLSA during postmortem ATP deprivation. Indeed, we also note that transient overexpression of some ALS-causing TDP-43 mutants (such as D169G and K181E) abolishes RNA-binding and phase separate into nuclear gels in cells in culture (*62*), but they ultimately cause classic cytoplasmic aggregates in patients’ nervous systems (*62*). Whether, and if so how, nuclear phase separation of TDP-43 correlates with and/or contributes to cytoplasmic aggregation is not yet determined. Nevertheless, our results suggest an essential partnership between TDP-43 and HSP70 chaperones in driving and maintaining TDP-43 phase separation. Future efforts are now needed to determine the identities of the TDP-43 acetylases and deacetylases and whether there are age-dependent changes in their activities that may synergize with the known age-dependent decline in proteasome activity (*63, 64*) to mediate stress-induced TDP-43 phase separation in age-related neurodegeneration.

## Acknowledgments

We thank Drs. E. Griffis and D. Bindels from the Nikon Imaging Center at UCSD for help on DIC and fluorescence microscopy. We acknowledge personnel at UCSD School of Medicine: Y. Jones and V. Taupin from the EM facility for sample preparation; Dr. J. Santini from the microscopy core for assistance in fluorescent microscopy. We thank Dr. T. Cohen for sharing TDP-43^2KQ^ variants.

## Funding

The authors thank the following funding support: H.Y., NIA AG059358 and NINDS NS114162; K.G., NSF DMS-1816630; D.S. Damon Runyon Foundation; J.M.N., the NSERC (RGPIN-2019-06435, RGPAS-2019-00014, DGECR-2019-00321), and NSF (DMS-171474, DMS-1816630); M.L. CIBERNED, CB06/05/0037; A.S.G., NIGMS GM081506; D.W.C., NINDS NS027036, the NOMIS Foundation; UCSD microscopy core, NINDS NS047101.

## Author contributions

H.Y. and D.W.C. designed most of the experiments. H.Y. performed most of the experiments. S.L. performed the proximity labeling and quantitative proteomic analyses. K.G. and J.N. designed and performed the mathematical modeling. D.S. and E.V. designed and performed the Cryo-ET. O.T. and M.L. designed and performed the EM and immunoEM in rat DRG neurons. D.T. and S.V-S. helped in generating key materials. J.R.Y. supervised the proteomic analyses. H.Y., K.G. and D.W.C. wrote the original draft. A.S.G., E.V., N.J., M.L., S.D.C. reviewed and edited the manuscript.

## Competing interests

Authors declare no competing interests.

## Data and materials availability

All data is available in the main text or the supplementary materials. Plasmids and cell lines are available upon request.

## Supplementary Materials

### Materials and Methods

#### Plasmids

Plasmid information is listed in Supplementary table S3. In brief, the entry vector (pHY135 and pHY195) were used to generate most of the lentiviral vectors. Traditional cloning method (using double-restriction digestion) and Gibson assembly were used. Most of mammalian expression vectors (not lentiviral vector) were directly ordered from Addgene. All plasmids constructed in this manuscript will be deposit to Addgene. TDP-43^2KQ^, and TDP-43^mNLS,2KQ^ cDNA was a gift from Dr. Todd Cohen. All other TDP-43 variants were constructed by H.Y. with the help of D. T. and S. L.

#### Cell culture, transfection and infection

Cell lines used in this paper are: HEK293T (ATCC: CRL-11268), U2OS (ATCC: HTB-96), SH-SY5Y (ATCC: CRL-2266), and human iPS cell line CV-C is a gift from Eugene Yeo’s group at UCSD (maintenance and differentiation will be described in a different session). Routine maintenance of these model cell lines follows the guideline posted on ATCC. In brief, U2OS and HEK293T cells were cultured in complete DMEM supplemented with 10% Fetal bovine serum. SH-SY5Y cells were cultured in DMEM/F12 supplemented with 10% Fetal bovine serum.

Transient transfection was performed for HEK293T cells and U2OS cells at 60% confluency, by using transfection reagent TransIT X2 (Mirus, MIR6000) and following standard protocol posted on the product page of TransIT X2. In brief, serum free DMEM was used to dissolve DNA, and then TransIT X2 was added to the mix. After 15 minutes, the final mix was added dropwise to attached cells. The culture medium was changed after 12~24 hours post transfection.

To package lentivirus, the 2nd generation packaging system was used. Briefly, 0.5 x106 per well of 293T cells were seeded in a 6-well plate. For lentiviral transfection, 2.5 μg of the lentiviral plasmid (constructs are labelled as lentivirus in Key Resources Table), 1.25 μg of pMD2.G and 0.625 μg of psPAX2 were inoculated to each well using Mirus transIT-X2 transfection reagent (Mirus). Culture medium was changed to fresh medium at 4~24 hours post transfection. Two days after transfection, the culture medium was filtered through a 0.45 μm syringe filter to generate the viral soup. The viral soup containing 10~50 μg/mL protamine sulfate was added to U2OS or SH-SY5Y cells for infection. The viral soup was removed 24 hours after infection, and cells were passaged at least once before selection. Infected cells are selected based on the selection marker encoded by the lentivirus. For U2OS, the concentrations of the antibiotics used for selection were 200 μg/mL for neomycin (Thermo Fisher), 20 μg/mL for blasticidin (Thermo Fisher), 1 μg/mL for puromycin (Thermo Fisher). For SH-SY5Y cells, the concentrations were 400 μg/mL for neomycin, 10 μg/mL for blasticidin and 3 μg/mL for puromycin. Detailed guides and protocols posted can be found on the Addgene website:

https://www.addgene.org/protocols/lentivirus-production/

https://www.addgene.org/guides/lentivirus/

#### Human induced pluripotent stem cell (iPSC) culture

The human iPSCs was grown in Matrigel-coated dishes (Corning,cat.# 354234), in mTeSR Plus media (StemCell Technologies, 05825) and passaged using enzyme-free dissociation medium (PBS based, EMD Millipore, S-014-B) or Accutase (Innovative Cell Technologies, Inc., AT-104). ROCK inhibitor Y27632 (Apexbio, B1293-10) was used at a concentration of 10μM for 24h after every passaging or thawing.

#### Motor neuron differentiation from human iPSCs

The iPSCs were differentiated according to a modified version of Martinez *et al.(62),* based on the established strategy of dual-SMAD inhibition (*63, 64*). iPSC dishes at 60-80% confluency were placed in **0.5XN2B27 media** (DMEM/F12 + GlutaMAX media (Life Technologies, 10565-018) supplemented with 0.5X N2 supplement (Life Technologies, 17502-048), 0.5X B27 supplement (Life Technologies, 17504-044), 0.1mM Ascorbic Acid (Sigma-Aldrich, A4544), and 1% PenStrep (Life Technologies, 15140-122)), which was supplemented with different drugs throughout differentiation. On days 1-6, cells were fed daily with 0.5XN2B27 media in the presence of 1μM Dorsomorphin (Tocris, 3093), 10μM SB431542 (Tocris, 1614) and 3μM CHIR99021 (Tocris, 4423). On day 7-18, cells were fed daily with 0.5X N2B27 in the presence of 1μM Dorsomorphin, 10μM SB431542, 0.2μM Smoothened Agonist (SAG) (EMD Biosciences, 566660) and 1.5μM Retinoic Acid (Sigma-Aldrich, R2625). On days 18-21, cells were fed every other day with 0.5XN2B27 in the presence of 0.2μM SAG and 1.5μM Retinoic Acid. On day 22 cells were passaged onto poly-Ornithine (Sigma, P3655) / poly-D-Lysine (Sigma, P6407) /Laminin (Sigma, L2020) coated plates, and on days 22-25 were fed with 0.5XN2B27 supplemented with 2ng/μl of BDNF(R&D Systems, 248-BD), CTNF(R&D Systems, 257NT) and GDNF (R&D Systems, 212GD), in the presence of 2μM DAPT (Tocris, 2634). From days 25 on, cells are fed every 2-3 days with 0.5XN2B27 supplemented with BDNF, CTNF and GDNF. Cells were treated with 5μM Ara-C (Sigma, C1768) for 48h to avoid expansion of glial cells. Cells were analyzed by immunofluorescence at different timepoints for the presence of ChAT, MAP2 and NFH (data not shown).

#### Fluorescence activated cell sorting (FACS)

Before cell sorting or flowcytometry analysis, cells were trypsinized and resuspended in FACS buffer (1% FBS in PBS). Cell suspension was passed 40 μm strainer (Corning #352235) to form single cell suspension. The cells were loaded to SONY SH800S cell sorter to sort for single cell clones, or a population of defined Clover fluorescence. For flowcytometry analyses, BD LSRFortessa was used.

#### Chemicals and cell treatments

Doxycycline (DOX Sigma-Aldrich, Cat# D9891) was used to induce gene expression via the TetON3G promoter. U2OS cell and SH-SY5Y cells were treated at 500ng/mL final concentration of DOX. Trichostatin A (TSA, ApexBio, Cat# A8183) at 10ug/mL and Vorinostat (SAHA, ApexBio, A4084) at 20μM final concentrations were used to inhibit HDAC activity in HEK293T, U2OS and SH-SY5Y cells. Bortezomib (BTZ, ApexBio, Cat# A2614) at 2.5μM final concentration was used to inhibit the proteasome function in SH-SY5Y cells and iPSC-derived motor neurons. RNA labeling was conducted by following the protocol of the kit (ThermoFisher, Cat# C10330). HSP70 inhibitor (VER155008, ApexBio, Cat# A4387) was used at 50μM final concentration.

#### Antibodies

Rabbit anti-TDP-43 antibody (Proteintech 12892-1-AP), mouse anti-GAPDH monoclonal antibody (Clone 6C5, Millipore CB1001) were used for immunostaining and immunoblotting. Goat anti-FUS (Bethyl A303-839A), Chicken anti-MAP2 (Novus NB300-213). Mouse anti-XFP (Clontech JL-8) and rabbit anti-TDP-43 (Proteintech 10782-2-AP) were used for immunoEM.

#### Fluorescence microscopy

Immunofluorescence and live fluorescence imaging were taken by using Zeiss LSM 880 confocal microscope with Airyscan or Leica SP8 confocal microscope with lightening deconvolution. Both scopes are at the microscopy core in the Department of Neurosciences. Before imaging, cells were seeded in glass-bottom 8-well μ-slide (iBidi 80827). For live imaging, cells were incubated in Leibovitz’s L-15 Medium supplemented with 10% FBS. For immunofluorescence, cells were first fixed with 4% PFA for 10 minutes at room temperature (RT), washed twice with PBS, and permeabilized by using 0.5% Triton X-100 in PBS. Then cells were incubated with blocking buffer (2% BSA, 0.1% Triton X-100 in PBS) for 30 minutes at RT. Primary antibodies were diluted in the blocking buffer, and cells were incubated with primary antibody over night at 4°C. Then cells were washed 3 times at RT by the washing buffer (0.1% Triton X-100 in PBS). Secondary antibodies were also diluted in the blocking buffer. Cells were incubated with secondary antibody for 45 minutes at RT, covered from light. After the incubation with secondary antibodies, cells were washed 4 times at RT by the washing buffer and stained with DAPI (Tocris #5748) at 1μg/mL in PBS. After DAPI staining, cells can be post-fixed by 4% PFA for 5 minutes at RT, and washed with PBS twice afterward, to immobilize the antibody on the antigen. Then cells were mounted with ProLong Gold (ThermoFisher P10144) and 8mm coverslips (Electron Microscopy Sciences #72296-08). The slides were cured at RT overnight in dark.

#### Differential interference contrast (DIC) microscopy and Complete Extinction microscopy (CEM)

Both DIC and CEM were performed by using Nikon Eclipse Ti2-E microscope, equipped with 100x lense, TC-C-DICP-I intelligent linear polarizer (adjustable), a build-in linear polarizer in the base unit, CO_2_ and temperature was maintained by Okolab H101 water jacket stage top chamber with lense heater, Okolab CO2 unit, and Oko-touch digital recorder. The sample temperature was recorded every 30s during the experiment. For DIC microscopy, the Normarski prism was inserted to the base of the lens, and the adjustable polarizer was turned to obtain optimal contrast. For CEM, the Normarski prism was pulled out, and the microscope was set to a position where only two polarizers are in the light pass. Images and movies were taken when the two linear polarizers are aligned at 0°, which allows maximum amount of light to pass. Then the adjustable linear polarizer was carefully turned to a position which the least amount of light was detected by the camera. This position is then defined as 90° (perpendicular). Images and movies were then taken at this position.

#### Fluorescence recovery after photobleaching (FRAP) analysis and quantification

FRAP experiments on U2OS cells were performed on Zeiss LSM880 Airyscan microscope with 40x/1.2 W objective. The intensity of the fluorescent signal is controlled in the detection range through changing the laser power, digital gain and off-set. For the green channel, bleaching was conducted by a 488-nm line from an argon laser at ~80%-100% intensity with ~10-20 iterations. FRAP experiments on SH-SY5Y TDP-43EGFP cells were performed on Olympus FV1000 Spectral Confocal using SIM Scanner with 100x oil immersion objective, bleaching was conducted scanning a region of 1×1 μm for 300ms at 8% of laser intensity at 405-nm. Fluorescence recovery was monitored at 0.5 seconds, 2 seconds, 5 seconds or 10 seconds intervals for 3 minutes. In the focal-bleach experiment roughly half of a particle is bleached or fully photobleached, and then the distribution of the fluorescence within the photo-manipulated particle is determined over time. During the experiment cells were maintained in Leibovitz’s L-15 medium (CO2 independent).

The FRAP data was quantified using Image J. The time series of the granule fluorescence intensity was calculated and the intensity of the background (area with no cells) was subtracted from the granule intensity. The intensity of the granule during the whole experiment was normalized to 1 before bleaching. An average of at least 10-20 particles per condition was used to calculate the mean and standard error. The averaged relative intensity and standard error were plotted to calculate the dynamics of the particles.

#### Immunoblotting

Sample collection: Cells were washed with cold PBS on ice before lysis. RIPA buffer was used to lyse cells. To remove DNA and RNA, and inhibit protease activity, RIPA buffer were supplemented with benzonase (Millipore #101654) at 1U/mL and protease inhibitor cocktail at 1X (100X cocktail from ThermoFisher #78430). Cells were lysed with 200uL per well for 6-well plate for 20 minutes and collected in 1.5mL tubes. Lysates were centrifuged at 15000g for 10 minutes. Supernatant were mixed with NuPAGE LDS sample buffer (ThermoFisher NP0007). All procedures were performed on ice or at 4°C before boiling. SDS page, transfer, and blotting procedures were performed by following standard protocol. ECL substrates were used to visualize HRP-conjugated secondary antibody on the blot, and Biorad ChemiDoc system was used to capture the luminescent signals.

#### Electron microscopy (EM) and ImmunoEM

U2OS cells were plated on coverslips and induced by doxycycline to express TDP-43 variants. Cells were fixed by 2% glutaraldehyde and in 0.1M sodium cacodylate (SC buffer) buffer for 60min or longer at 4°C. After fixation, all procedures are carried out on ice. Fixed cells were washed with 0.1M SC buffer five times, and stainded with 1% OsO_4_ in 0.1M SC buffer for 45 minutes. Stained cells were then washed with 0.1M SC buffer five times and MilliQ water for 2 times. Coverslips were placed in 2% uranyl acetate buffer for 45 minutes and then rinced with Milli-Q water. Samples were serial dehydrated by 20%, 50%, 70%, 90%, and 2 times in 100% ETOH for 1 minute for each time and then dry acetonefor 2X1mintue. After dehydration, all steps were carried at room temperature (RT). Samples were then incubated at with Ducurpan:Acetone=50/50 for 1 hour, and then two times of 1 hour incubation in fresh 100% Durcupan. Coverslips were embedded in Durcupan resin, and cuted into 60nm ultrathin sections by diamond knife. Sections were mounted on 300 mesh grids.

For immunoEM studies, U2OS cells containing TDP-43 intranuclear spherical annuli were fixed for 12h in 4% PFA in 0.1 M phosphate buffer, pelleted in 10% gelatin, cryoprotected in sucrose, and snap frozen in liquid nitrogen. Ultrathin cryosections (70–80 nm) were cut as previously described (Zheng et al., 2004). Sections were picked up with a 1:1 mixture of 2.3 M sucrose and 2% methyl cellulose (15cp) as described by Liou et al. (1996) and transferred onto Formvar and carbon-coated copper grids. Immunolabeling was performed by a slight modification of the “Tokuyasu technique”, (Tokuyasu, 1980). Briefly, grids were placed on 2% gelatin at 37°C for 20 minutes, rinsed with 0.15 M glycine/PBS and the sections were blocked using 1% cold water fish-skin gelatin. For immunogold labeling of TDP-43^2KQ-clover^, sections were sequentially incubated for 2h with a mouse anti-GFP antibody (Clontech Living Colors A.V. JL-8), followed by a 12nM gold conjugated goat anti-mouse IgG (Jackson Immunoresearch, 115-205-071) for 1h. Grids were then contrasted for 10 min in 0.4% uranyl acetate and 1.8% methyl cellulose on ice.

For immunoEM of TDP-43 in rat tissue, bortezomib-treated rats were subjected to transcardial perfusion under deep anesthesia with 3.7 % paraformaldehyde in 0.1 M cacodylate buffer for 15 min at RT. Small tissue fragments of sensory ganglia were dissected and post-fixed in the same fixative solution for 4h at RT, washed in 0.1 M cacodylate buffer, dehydrated in increasing concentrations of methanol at −20 °C, embedded in Lowicryl K4M at −20 °C, and polymerized with ultraviolet irradiation. Ultrathin sections were mounted on Formvar coated nickel grids and sequentially incubated with 0.1 M glycine in PBS for 15 min, 3 % BSA in PBS for 30 min, and the primary rabbit polyclonal anti-TDP-43 antibody (diluted 1:100 in 50 mM Tris-HCl, pH 7.6, containing 1 % BSA and 0.1 M glycine) for 1 h at 37 °C. After washing, sections were incubated with the goat anti-mouse IgG antibody coupled to 10-nm gold particles (BioCell, UK; diluted 1:50 in PBS containing 1 % BSA). Following immunogold labeling, the grids were stained with lead citrate and 2% (w/v) uranyl acetate. As controls, ultrathin sections were treated as described above but with the primary antibody being omitted.

Imaging was carried out using a JEOL 1200 EX II electron microscope equipped with an Orius CCD Gatan camera and Gatan digital micrograph software (Gatan, Pleasanton, CA; University of California, San Diego, Cellular and Molecular Medicine Electron Microscopy Facility).

#### Cryo-FIB milling and cryo-electron tomography

U2OS cells were dox-induced for 2-6 days and then seeded on glow-discharged Quantifoil grids (R1/4, Au 200-mesh grid, Electron Microscopy Sciences). The cells were then cultured for 24 hrs with continued dox-induction. The grids were then manually blotted from the backside only (to remove excess media) and quickly plunge-frozen into liquid ethane/propane mixture using the custom-built vitrification device (Max Planck Institute for Biochemistry, Munich). From thereon, grids were always kept in cryogenic conditions and utmost care was taken to prevent ice contamination. The grids were then clipped onto FIB-AutoGrids (ThermoFisher) for milling inside a Thermo Scientific Aquilos DualBeam. ~120-150 nm lamellae (~11 mm in width) were prepared in the Aquilos using rectangular milling patterns as previously described^4,5^. Briefly, the coarse milling was performed with the ion beam current of 0.10-0.50 nA. The current was reduced to 10-50 pA for careful and slow fine milling.

The lamellae created on the grids were then stored in cryogenic conditions and eventually transferred to the transmission electron microscope (Titan Krios G3; ThermoFisher) using the AutoLoader (ThermoFisher). Tilt-images, in the form of cryo-transmission electron images, were acquired on the microscope operated at 300 kV and equipped with post-column energy filter (Gatan). The images were recorded on a K2 Summit (Gatan) direct detector in the counting mode with the help of SerialEM software. The images were recorded in the movie mode. The tilt-series acquisition parameters were as follows: tilt range: ± 60, magnification: 26000X-19500X, tilt increment: 3/2/1.5°, pixel size of 0.53-0.71 nm, target defocus: −5 μm. The frames of the tilt-series images were motion corrected and dose-weighted using motioncorr2. The tilt-series images were then aligned in IMOD using patch-tracking. No CTF correction was performed. The weighted back-projection method was used for the final tomographic reconstruction.

#### Proximity labeling and enrichment of biotinylated protein

Before the labeling experiment, U2OS cells were passaged in (Lys0, Arg0) or heavy (Lys8, Arg10) DMEM medium for 5 generations. U2OS cells were incubated with 500 μM biotin phenol (Iris-Biotech, 41994-02-9) containing medium for 30 min. Then 1 mM hydrogen peroxide was added to the medium to activate APEX reaction for 1 min, followed by immediate quenching of reaction with ice-cold quenching buffer (1xPBS, 10 mM sodium azide, 10 mM sodium ascorbate, 5 mM Trolox). After wash with cold quenching buffer for three times, the cells were collected from plates with scrapers.

Cells were lysed in lysis buffer (100 mM NaPO4, PH 8.0, 8 M Urea, 0.05% SDS, 10 mM sodium azide, 10 mM sodium ascorbate, 5 mM Trolox supplemented with 500 unit benzonase to remove DNA and RNA) and rotate at room temperature for 15 min. Then increase the concentration of SDS to 1% and sonicate at water bath sonicator for 10 mins. Protein concentration was measured using 2-D quant kit (GE healthcare, Cat# 80648356), by following manufacturer’ instruction. The denatured proteins were reduced with 5 mM TCEP and alkylated with 10 mM Iodoacetamide. Equal amount of proteins from light and heavy group were mixed together and then the mixed cell lysates were diluted with equal volume of ddH2O to reduce the concentration of urea to 4 M and SDS to 0.5%. The samples were incubated with streptavidin magnetic beads at 4°C overnight. After four washes with wash buffer (100 mM NaPO4, PH 8.0, 4 M Urea), the beads were resuspended in 100 mM TEAB, 2 M Urea supplemented with 10 ng/uL Trypsin, 5 ng/uL Lys-C for digestion overnight. The digested products were collected and added with 1% formic acid (final concentration) for LC-MS/MS analysis.

#### Liquid chromatography-Mass spectrometry analysis

LC-MS/MS was conducted on Orbitrap Elite Hybrid Mass Spectrometer interfaced with a nano-flow HPLC Easy II. Digested peptides were loaded onto a 100 μm x 15 cm analytical column packed with 3 μm, 120 Å C18 resin (). The peptides were separated over a 90-min linear gradient from 0% buffer B (95% acetonitrile, 0.1% formic acid), 100% buffer A (5% acetonitrile, 0.1% formic acid) to 30% buffer B, followed by a 10-min gradient from 27% to 80% buffer B, then maintaining at 80% buffer B for 20 min. The flow rate was 300 nL/min. The MS parameters were: R = 140,000 in full scan, R = 7,500 in HCD MS2 scan; the 10 most intense ions in each full scan were selected for HCD dissociation; the AGC targets were 1e6 for FTMS full scan and 5e4 for MS2; minimal signal threshold for MS2 was 5e3; precursors having a charge state of +1, or unassigned were excluded; normalized collision energy was set to 30; dynamic exclusion was on.

#### Quantitative mass spectrometry data analysis

The raw data was processed by pParse (*65*), and co-eluting precursor ions were excluded. MS1 and MS2 data were searched using pFind 3 (*66*) against a complete human protein database downloaded from Uniprot with the addition of APEX2 and Clover protein sequence. The settings of pFind 3 are: open search mode; 3 missed cleavage sites for trypsin digestion; peptide length 6-25 aa; fixed modification was carboxymethylation on Cys (58.0054 Da); variable modifications were oxidation on Met (15.9949 Da) and acetylation on N-term (42.0105 Da). The search results are filtered with less than 10 ppm mass deviation for precursor ions and 20 ppm for fragment ions, FDR < 0.01 at peptide level. The identified spectra were used for quantification with pQuant (*67*) with the setting of using 200 MS1 scans before and after the spectra corresponding to identified spectra for searching the chromatography peaks and allowing 2 holes in the chromatography peak and the intensity of ion within 10 ppm mass deviation was used.

The median value of the ratio of light to heavy peptides from each protein was used as the ratio of the protein. The ratios of each protein from two forward labeling groups and one reverse labeling group were used to calculate P-value through one sample t-test. The volcano plot was generated with R package.

#### Statistics

Except the quantitative analyses for the proteomic data, student T-test was used to calculate significance, and P<0.05 is considered statistically significant.

## References

1. S. F. Banani, H. O. Lee, A. A. Hyman, M. K. Rosen, Biomolecular condensates: organizers of cellular biochemistry. Nat Rev Mol Cell Biol 18, 285–298 (2017).

2. W. T. Snead, A. S. Gladfelter, The Control Centers of Biomolecular Phase Separation: How Membrane Surfaces, PTMs, and Active Processes Regulate Condensation. Mol Cell 76, 295–305 (2019).

3. G. Valentin, Repertorium fur Anatomie und Physiologie. MVerlag Veit Comp. Berl. 1, 1–293 (1836).

4. R. Wagner, Einige Bemerkungern und Fragen uber das Keimblaschen. Mullers Archiv Anat. Physiol. Wiessenschaft Med., 373–377 (1835).

5. C. P. Brangwynne, T. J. Mitchison, A. A. Hyman, Active liquid-like behavior of nucleoli determines their size and shape in Xenopus laevis oocytes. Proc Natl Acad Sci U S A 108, 4334–4339 (2011).

6. C. P. Brangwynne et al., Germline P granules are liquid droplets that localize by controlled dissolution/condensation. Science 324, 1729–1732 (2009).

7. A. Jain, R. D. Vale, RNA phase transitions in repeat expansion disorders. Nature 546, 243–247 (2017).

8. J. B. Woodruff et al., The Centrosome Is a Selective Condensate that Nucleates Microtubules by Concentrating Tubulin. Cell 169, 1066–1077 e1010 (2017).

9. H. B. Schmidt, D. Gorlich, Transport Selectivity of Nuclear Pores, Phase Separation, and Membraneless Organelles. Trends Biochem Sci 41, 46–61 (2016).

10. D. Hnisz, K. Shrinivas, R. A. Young, A. K. Chakraborty, P. A. Sharp, A Phase Separation Model for Transcriptional Control. Cell 169, 13–23 (2017).

11. A. Patel et al., A Liquid-to-Solid Phase Transition of the ALS Protein FUS Accelerated by Disease Mutation. Cell 162, 1066–1077 (2015).

12. A. Molliex et al., Phase separation by low complexity domains promotes stress granule assembly and drives pathological fibrillization. Cell 163, 123–133 (2015).

13. D. Bracha et al., Mapping Local and Global Liquid Phase Behavior in Living Cells Using Photo-Oligomerizable Seeds. Cell 175, 1467–1480 e1413 (2018).

14. P. Li et al., Phase transitions in the assembly of multivalent signalling proteins. Nature 483, 336–340 (2012).

15. G. Pelzl, A. Hauser, Birefringence and phase transitions in liquid crystals. Phase Transitions 37, 33–62 (1991).

16. J. P. Taylor, R. H. Brown, Jr., D. W. Cleveland, Decoding ALS: from genes to mechanism. Nature 539, 197–206 (2016).

17. C. Amador-Ortiz et al., TDP-43 immunoreactivity in hippocampal sclerosis and Alzheimer’s disease. Annals of Neurology: Official Journal of the American Neurological Association and the Child Neurology Society 61, 435–445 (2007).

18. P. T. Nelson et al., Limbic-predominant age-related TDP-43 encephalopathy (LATE): consensus working group report. Brain 142, 1503–1527 (2019).

19. M. Polymenidou et al., Long pre-mRNA depletion and RNA missplicing contribute to neuronal vulnerability from loss of TDP-43. Nat Neurosci 14, 459–468 (2011).

20. E. Buratti, TDP-43 post-translational modifications in health and disease. Expert Opin Ther Targets 22, 279–293 (2018).

21. F. Gasset-Rosa et al., Cytoplasmic TDP-43 De-mixing Independent of Stress Granules Drives Inhibition of Nuclear Import, Loss of Nuclear TDP-43, and Cell Death. Neuron 102, 339–357 e337 (2019).

22. J. R. Klim et al., ALS-implicated protein TDP-43 sustains levels of STMN2, a mediator of motor neuron growth and repair. Nat Neurosci 22, 167–179 (2019).

23. J. P. Ling, O. Pletnikova, J. C. Troncoso, P. C. Wong, TDP-43 repression of nonconserved cryptic exons is compromised in ALS-FTD. Science 349, 650–655 (2015).

24. Z. Melamed et al., Premature polyadenylation-mediated loss of stathmin-2 is a hallmark of TDP-43-dependent neurodegeneration. Nat Neurosci 22, 180–190 (2019).

25. K. Miskiewicz et al., HDAC6 is a Bruchpilot deacetylase that facilitates neurotransmitter release. Cell Rep 8, 94–102 (2014).

26. A. Bhardwaj, M. P. Myers, E. Buratti, F. E. Baralle, Characterizing TDP-43 interaction with its RNA targets. Nucleic Acids Res 41, 5062–5074 (2013).

27. T. J. Cohen et al., An acetylation switch controls TDP-43 function and aggregation propensity. Nat Commun 6, 5845 (2015).

28. J. R. Mann et al., RNA Binding Antagonizes Neurotoxic Phase Transitions of TDP-43. Neuron 102, 321–338 e328 (2019).

29. Y. Chen, T. J. Cohen, Aggregation of the nucleic acid-binding protein TDP-43 occurs via distinct routes that are coordinated with stress granule formation. J Biol Chem 294, 3696–3706 (2019).

30. T. Afroz et al., Functional and dynamic polymerization of the ALS-linked protein TDP-43 antagonizes its pathologic aggregation. Nat Commun 8, 45 (2017).

31. A. Wang et al., A single N-terminal phosphomimic disrupts TDP-43 polymerization, phase separation, and RNA splicing. EMBO J 37, (2018).

32. N. Vo, R. H. Goodman, CREB-binding protein and p300 in transcriptional regulation. J Biol Chem 276, 13505–13508 (2001).

33. T. J. Cohen, V. M. Lee, J. Q. Trojanowski, TDP-43 functions and pathogenic mechanisms implicated in TDP-43 proteinopathies. Trends Mol Med 17, 659–667 (2011).

34. Y. M. Ayala et al., TDP-43 regulates its mRNA levels through a negative feedback loop. EMBO J 30, 277–288 (2011).

35. A. C. Elden et al., Ataxin-2 intermediate-length polyglutamine expansions are associated with increased risk for ALS. Nature 466, 1069–1075 (2010).

36. J. D. Vargas, E. M. Hatch, D. J. Anderson, M. W. Hetzer, Transient nuclear envelope rupturing during interphase in human cancer cells. Nucleus 3, 88–100 (2012).

37. S. Saha et al., Polar Positioning of Phase-Separated Liquid Compartments in Cells Regulated by an mRNA Competition Mechanism. Cell 166, 1572–1584 e1516 (2016).

38. R. Oldenbourg, Polarized light microscopy: principles and practice. Cold Spring Harb Protoc 2013, (2013).

39. C. Reyes Mateo, A. Ulises Acuna, J. C. Brochon, Liquid-crystalline phases of cholesterol/lipid bilayers as revealed by the fluorescence of trans-parinaric acid. Biophys J 68, 978–987 (1995).

40. K. Liu et al., Thermotropic liquid crystals from biomacromolecules. Proc Natl Acad Sci U S A 111, 18596–18600 (2014).

41. O. Rog, S. Kohler, A. F. Dernburg, The synaptonemal complex has liquid crystalline properties and spatially regulates meiotic recombination factors. Elife 6, (2017).

42. M. Mompean et al., Point mutations in the N-terminal domain of transactive response DNA-binding protein 43 kDa (TDP-43) compromise its stability, dimerization, and functions. J Biol Chem 292, 11992–12006 (2017).

43. Y. Shiina, K. Arima, H. Tabunoki, J. Satoh, TDP-43 dimerizes in human cells in culture. Cell Mol Neurobiol 30, 641–652 (2010).

44. K. Gasior et al., Partial demixing of RNA-protein complexes leads to intradroplet patterning in phase-separated biological condensates. Phys Rev E 99, 012411 (2019).

45. R. Rosenzweig, N. B. Nillegoda, M. P. Mayer, B. Bukau, The Hsp70 chaperone network. Nat Rev Mol Cell Biol 20, 665–680 (2019).

46. D. S. Williamson et al., Novel adenosine-derived inhibitors of 70 kDa heat shock protein, discovered through structure-based design. J Med Chem 52, 1510–1513 (2009).

47. M. Feric et al., Coexisting Liquid Phases Underlie Nucleolar Subcompartments. Cell 165, 1686–1697 (2016).

48. B. A. Gibson et al., Organization of Chromatin by Intrinsic and Regulated Phase Separation. Cell 179, 470–484 e421 (2019).

49. D. R. Palleros, W. J. Welch, A. L. Fink, Interaction of hsp70 with unfolded proteins: effects of temperature and nucleotides on the kinetics of binding. Proc Natl Acad Sci U S A 88, 5719–5723 (1991).

50. P. J. Lukavsky et al., Molecular basis of UG-rich RNA recognition by the human splicing factor TDP-43. Nat Struct Mol Biol 20, 1443–1449 (2013).

51. M. Ganassi et al., A Surveillance Function of the HSPB8-BAG3-HSP70 Chaperone Complex Ensures Stress Granule Integrity and Dynamism. Mol Cell 63, 796–810 (2016).

52. K. Illmensee, A. P. Mahowald, M. R. Loomis, The ontogeny of germ plasm during oogenesis in Drosophila. Dev Biol 49, 40–65 (1976).

53. K. E. Kistler et al., Phase transitioned nuclear Oskar promotes cell division of Drosophila primordial germ cells. Elife 7, (2018).

54. J. O. Narcis et al., Accumulation of poly(A) RNA in nuclear granules enriched in Sam68 in motor neurons from the SMNDelta7 mouse model of SMA. Sci Rep 8, 9646 (2018).

55. M. C. Gallego-Iradi et al., N-terminal sequences in matrin 3 mediate phase separation into droplet-like structures that recruit TDP43 variants lacking RNA binding elements. Lab Invest 99, 1030–1040 (2019).

56. S. C. Ling, M. Polymenidou, D. W. Cleveland, Converging mechanisms in ALS and FTD: disrupted RNA and protein homeostasis. Neuron 79, 416–438 (2013).

57. M. Neumann et al., Ubiquitinated TDP-43 in frontotemporal lobar degeneration and amyotrophic lateral sclerosis. Science 314, 130–133 (2006).

58. J. L. Robinson et al., Neurodegenerative disease concomitant proteinopathies are prevalent, age-related and APOE4-associated. Brain 141, 2181–2193 (2018).

59. H. S. Yang et al., Evaluation of TDP-43 proteinopathy and hippocampal sclerosis in relation to APOE epsilon4 haplotype status: a community-based cohort study. Lancet Neurol 17, 773–781 (2018).

60. K. A. Josephs et al., TDP-43 is a key player in the clinical features associated with Alzheimer’s disease. Acta Neuropathol 127, 811–824 (2014).

61. M. Neumann et al., TDP-43 in the ubiquitin pathology of frontotemporal dementia with VCP gene mutations. J Neuropathol Exp Neurol 66, 152–157 (2007).

62. H. J. Chen et al., RRM adjacent TARDBP mutations disrupt RNA binding and enhance TDP-43 proteinopathy. Brain 142, 3753–3770 (2019).

63. E. K. Sacramento et al., Reduced proteasome activity in the aging brain results in ribosome stoichiometry loss and aggregation. bioRxiv, 577478 (2020).

64. I. Saez, D. Vilchez, The Mechanistic Links Between Proteasome Activity, Aging and Age-related Diseases. Curr Genomics 15, 38–51 (2014).

