## Supplementary Figures and Legends for "TDP-43 and HSP70 phase separate into anisotropic, intranuclear liquid spherical annuli"

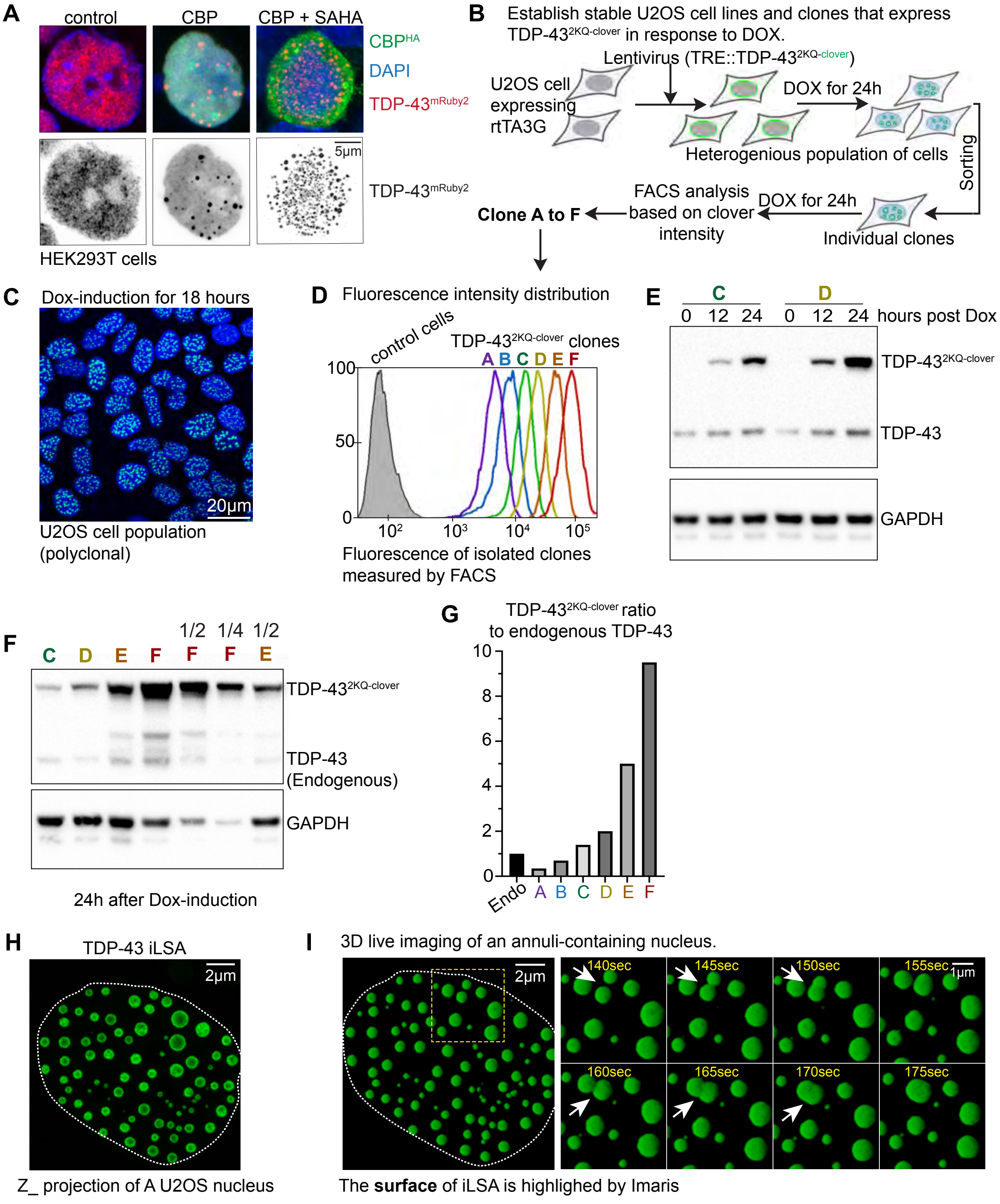


**Fig. S1. TDP-43 iLSA formation is driven by acetylation or expression of acetylation-mimic variant**

**Fig. S1. TDP-43 iLSA formation is driven by acetylation or expression of acetylation-mimic variant.** (**A**) Expressing acetyltransferase CBP promotes phase separation of nuclear TDP-43, which is further enhanced by the addition of pan-HDAC inhibitor SAHA. (**B**) A diagram explains the process of establishing U2OS cell lines and clones expressing different level of TDP-$43^{2KQ\text{-}clover}$. (**C**) Inducible expression of TDP-$43^{2KQ\text{-}clover}$ de-mixes in every cell. (**D**) Single U2OS cell clones (clone A to F) show dose-dependent expression of TDP-$43^{2KQ\text{-}clover}$ after 24 hours of DOX-induction. (**E**) Inducible expression of TDP-$43^{2KQ\text{-}clover}$ is time dependent. Immunoblot of TDP-43 is shown. (**F**) Immunoblotting of TDP-43 shows the relative expression level of TDP-$43^{2KQ\text{-}clover}$ to the endogenous TDP-43. Clone C, D, E and F were used because TDP-$43^{2KQ\text{-}clover}$ cannot be detected by immunoblotting in Clone A and B. (**G**) The expression level of TDP-$43^{2KQ\text{-}clover}$ relative to the endogenous TDP-43 based on the results of (**D**), (**E**) and (**F**). The geometric mean of each curve in (**D**) was used to calculate the relative expression, which matches the intensity measurements of immunoblotting (**F**). (**H, I**) Fluorescence 3D live-cell imaging reveals rapid fusion events (*Note: the 3D reconstruction highlights the surface of the iLSA, so the hollow center is masked*).

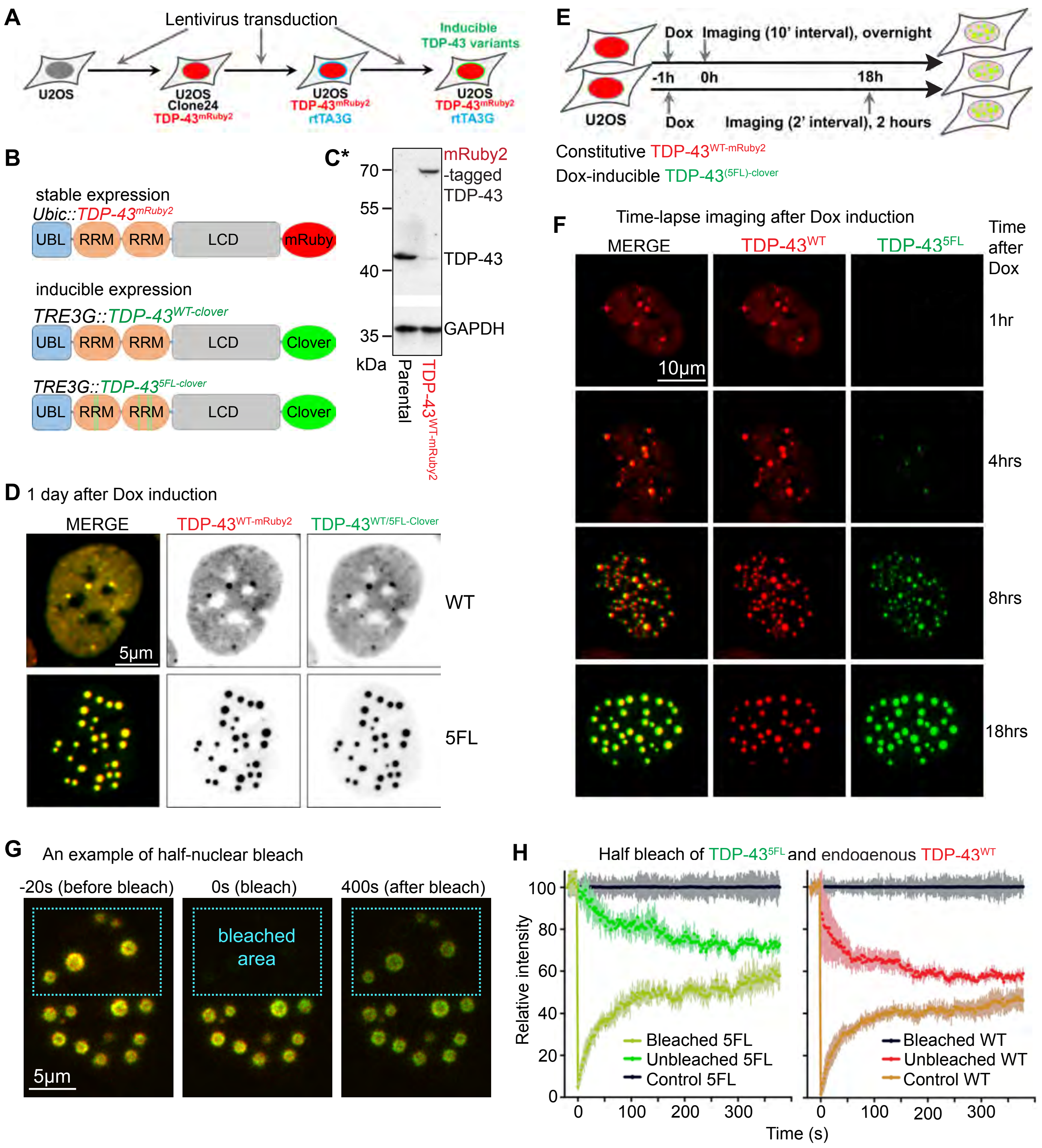


***NOTE: Panel C is adapted from Gasset-Rosa F et al, Supplementary Figure 1C**

Figure S2 RNA-binding deficient TDP-43 variant (TDP-435FL) de-mixes wildtype TDP-43 into iLSA

**Fig. S2. RNA-binding deficient TDP-43 variant (TDP-43$^{5FL}$) de-mixes wildtype TDP-43 into iLSA.** (**A**) A diagram explains the step-by-step process to establish a U2OS cell line that stably expresses TDP-43$^{WT\text{-}mRuby2}$ (red fluorescence) and inducible-expresses clover-tagged TDP-43 variants (green fluorescence). (**B**) The diagram of tagged TDP-43 wildtype or 5FL proteins. (**C**) Immunoblot shows TDP-43$^{WT\text{-}mRuby2}$ expresses at endogenous level. (**D**) Similar to TDP-43$^{2KQ\text{-}clover}$, inducible expression of TDP-43$^{5FL\text{-}clover}$ de-mixes TDP-43WT-mRuby2 into many iLSA (round droplets under low magnification), while inducible expression of TDP-43$^{WT\text{-}clover}$ only shows a handful of droplets. (**E**) and (**F**) Time-lapse imaging shows de-mixing of TDP-43$^{5FL\text{-}clover}$ is dominant over TDP-43$^{WT\text{-}mRuby2}$. (**G**) The dynamic intra-annuli molecular exchange is demonstrated by half-nuclear bleach of both the TDP-43$^{5FL\text{-}clover}$ and the TDP-43$^{WT\text{-}mRuby2}$. The FRAP recovery curve is shown in (**H**).

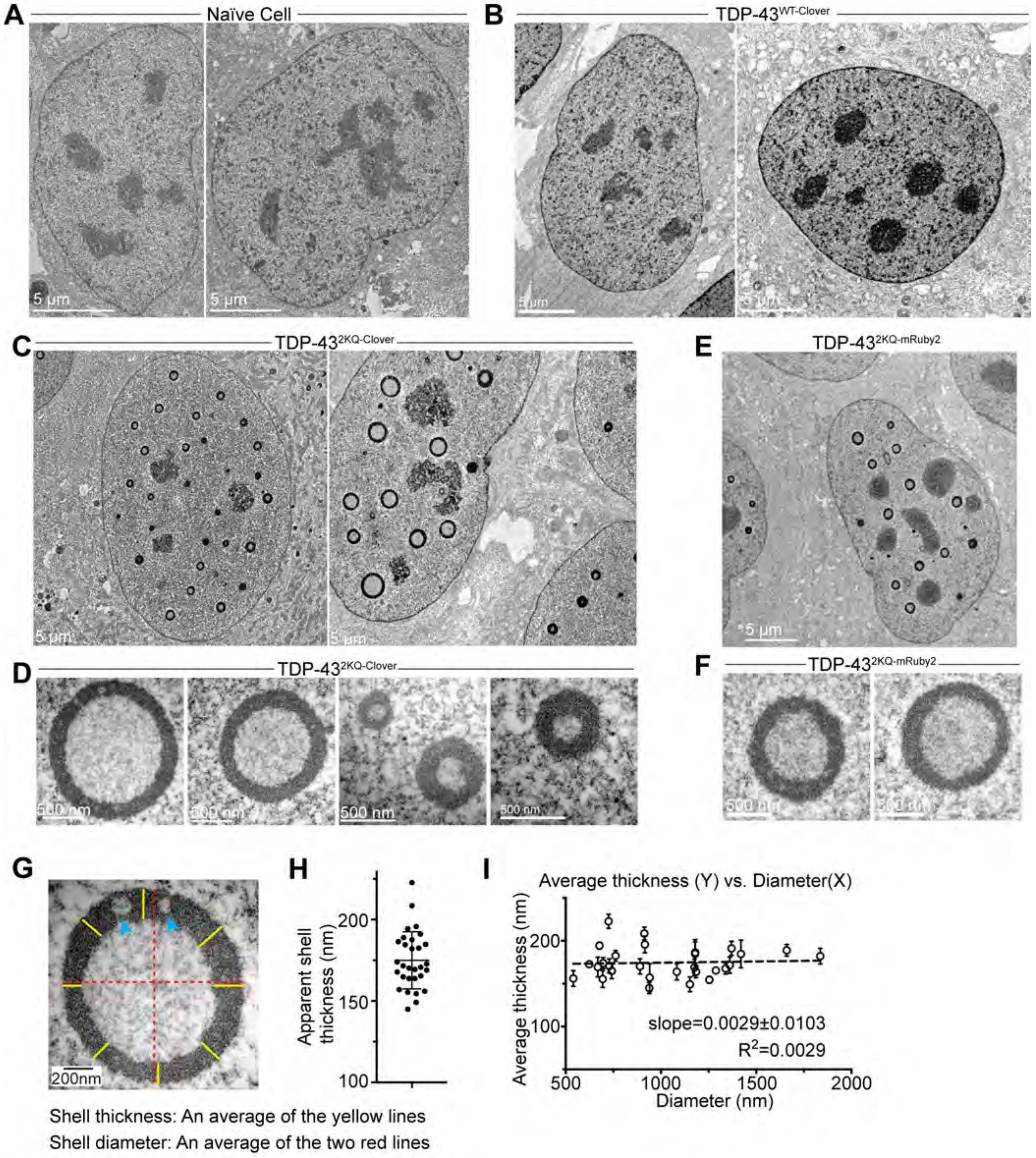


**Fig. S3. TDP-43 iLSA is directly observed by TEM.**

**Fig. S3. TDP-43 iLSA is directly observed by TEM.** (**A**) Two more examples of nuclei of naïve U2OS cells. (**B**) Two more examples of nuclei of U2OS cells expressing TDP-43$^{WT\text{-}clover}$. (**C**) Two more examples of nuclei of U2OS cells expressing TDP-43$^{2KQ\text{-}clover}$. (**D**) Magnified examples of big and small TDP-43$^{2KQ\text{-}clover}$ iLSA. (**E**) TDP-43$^{2KQ\text{-}mRuby2}$ also forms iLSA. (**F**) TDP-43$^{2KQ\text{-}mRuby2}$ iLSA show no difference to TDP-43$^{2KQ\text{-}clover}$ iLSA under TEM. Two magnified examples are shown (**G**) An example shows how the thickness and diameter of annuli were measured. The average of the red lines is the diameter of the annulus, while the average of the yellow lines is the thickness of the annulus. Blue arrows point to two electron-lucent regions, which may be putative fusion events or material exchanged events. (**H**) The average thickness plot of the annuli. (**I**) The thickness of the iLSA remains constant. Annuli were plotted by values of the average thickness (Y-axis) and the diameter (X-axis). Linear regression calculated the slope of the dashed line, which is close to 0, and $R^2$=0.029 shows no correlation between the thickness and the diameter.

A

B

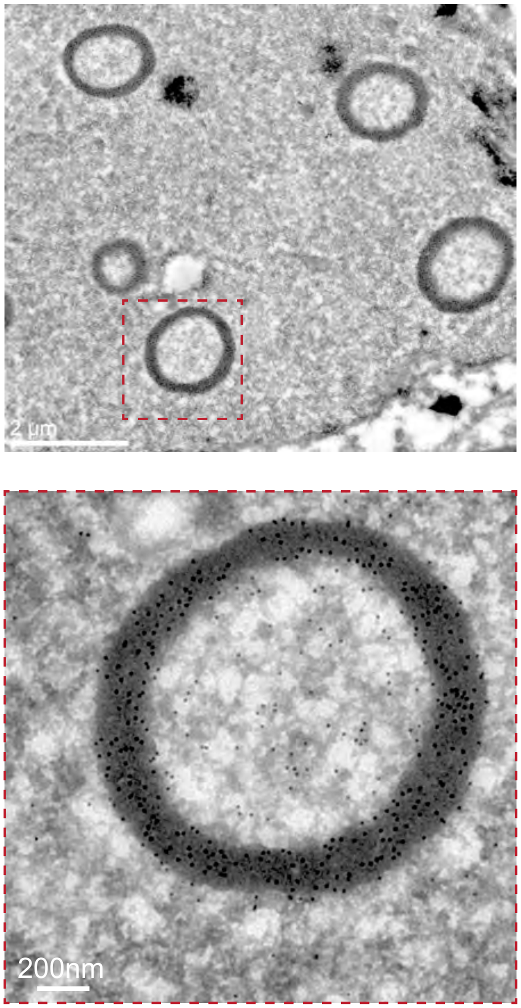


**Fig. S4. Immunogold labeling of U2OS cells expressing TDP-435FL-clover.** (A) An overview of a cross-section of a nucleus, where shows several iLSA are stained with anti-GFP gold nanoparticles. The highlighted annuli (red box) is magnified in panel (B) to show the distribution of gold nanoparticles.

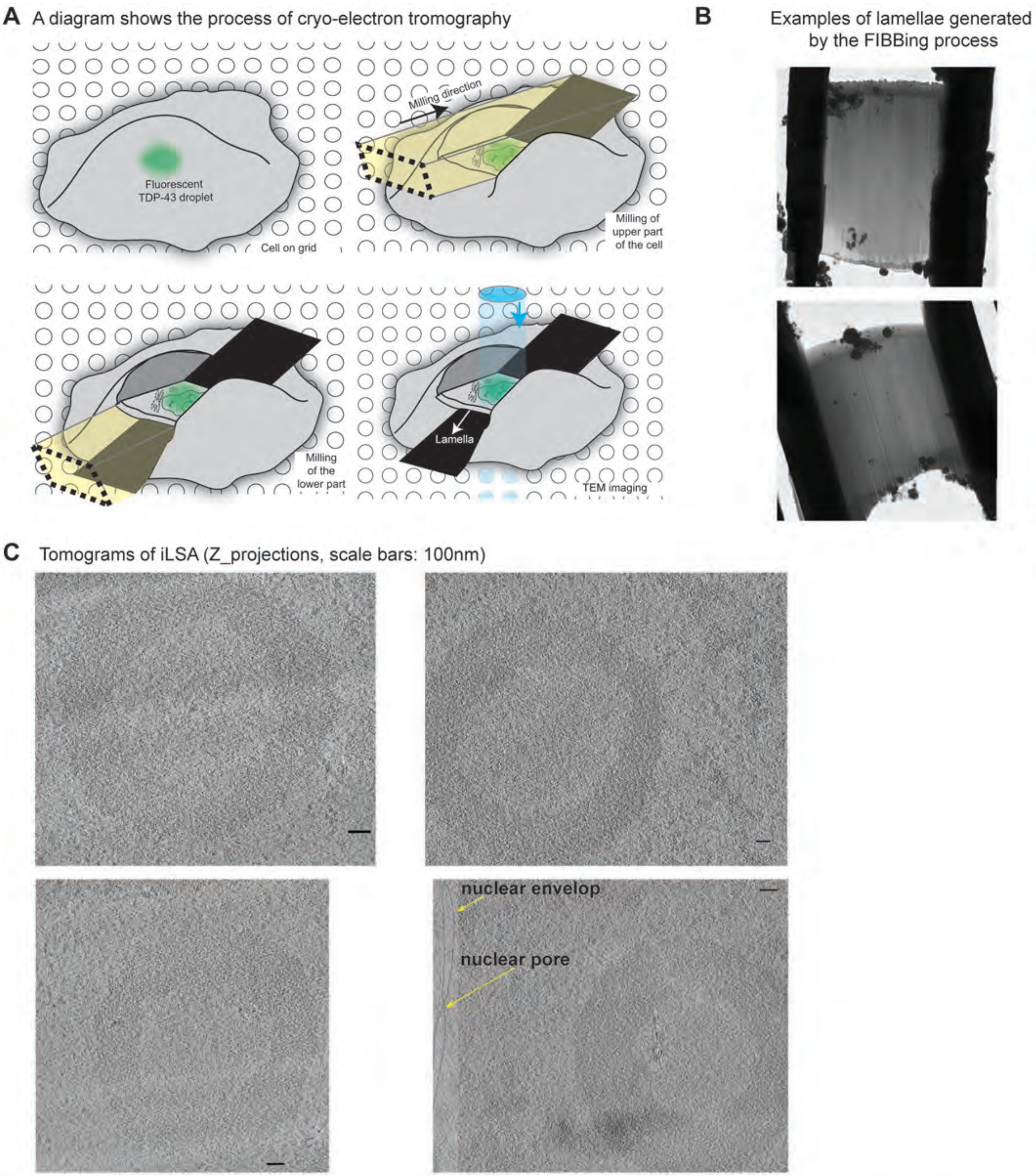


**Fig. S5. Frozen hydrated annuli are observed *in situ* by cryo-electron tomography (CryoET).** (A) A diagram explains the workflow of CryoET. Cells were seeded on plasma-treated carbon grids and then snap-frozen. A lamella was processed by focus ion beam. The lamella was imaged by TEM. (B) Lamellae were used to acquire CryoET images of annuli. (C) More images of TDP-43 annuli resolved by CryoET (scale bar: 100nm).

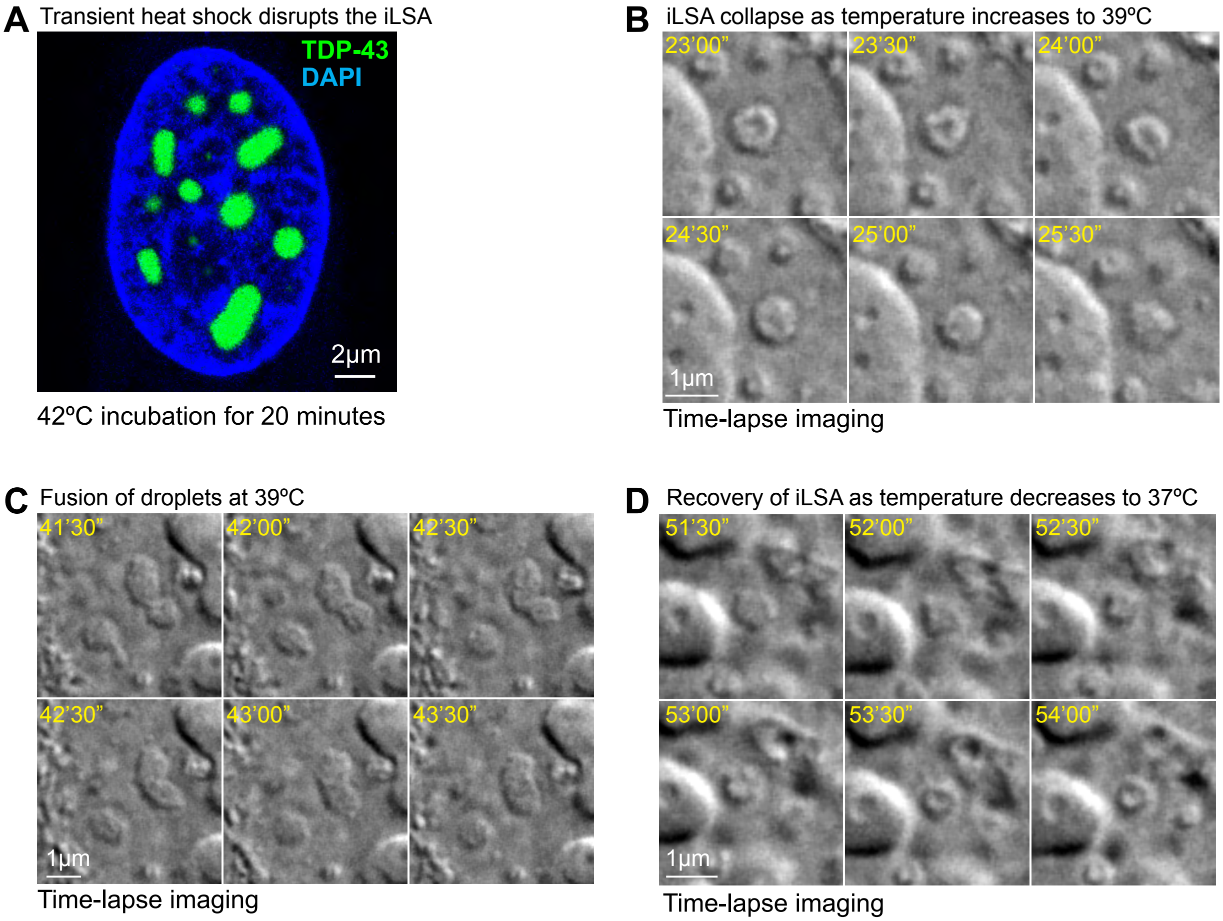


**Fig. S6. High physiological temperature disrupts TDP-43 iLSA.** (A) A fluorescent image shows TDP-43 annuli collapsed to oval droplets after 30 minutes of heat shock at 42ºC. (B) to (D) Time lapse imaging with better time resolution, correlates with Fig. 4I-K.

k

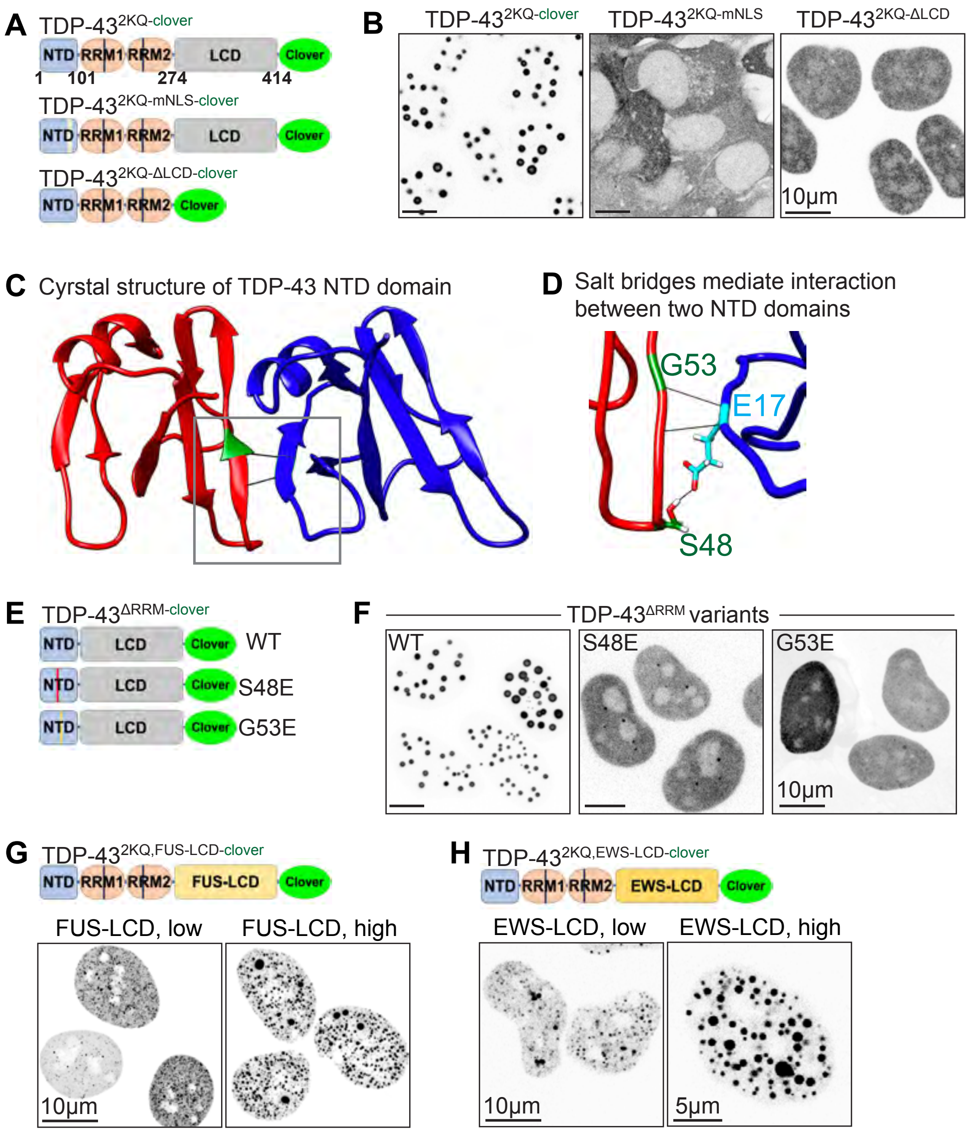


**Fig. S7. Both the N-terminal oligomerization domain and the C-terminal LCD domain are required for iLSA formation.** (A) A diagram of inducible-expressed TDP-43 variants (TDP-432KQ-clover, TDP-432KQ,mNLS-clover and TDP-432KQ,ΔLCD-clover) in U2OS cells. (B) Monochrome images show the distribution of different TDP-43 variants showed in the panel A. Mutation in the NLS or deletion of the LCD domain abolishes annular phase separation. (C) A diagram of inducible-expressed TDP-43 variants (TDP-43 ΔRRM-clover, TDP-43Δ RRM,S48E-clover and TDP-43ΔRRM,G53E-clover) in U2OS cells. (D) The structure of the NTD dimer shows salt bridges formed between G53/S48 of one molecule and E17 from the other. (E) Monochrome images show the distribution of different TDP-43 variants showed in the panel D. Removing the RNA-binding domains (ΔRRM) also drove annular phase separation. However, single amino acid substitutions (S48E or G53E) that disrupted the NTD dimer interface completely abolished phase separation. (F) A diagram of inducible-expressed TDP-43 variants with LCD-swap (TDP-432KQ,FUS-LCD-clover and TDP-432KQ,EWS-LCD-clover) in U2OS cells. (G) LCD swap mutants form intranuclear droplets but not spherical annuli. Low and high expression cells are shown for both FUS-LCD and EWS-LCD swap mutants.

**A** Demixing of RNA and TDP-43$^{2KQ}$ only produces uniform droplets

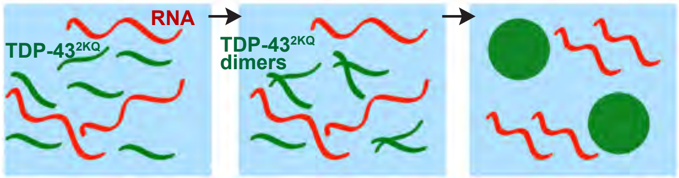


**B** Equations for Model of demixing including RNA and TDP-43$^{2KQ}$

$$\frac{\partial P}{\partial t} = \nabla \cdot [\lambda_P \nabla P] - 2k_{on} P \cdot P + 2k_{off} P_2$$

$$\frac{\partial P_2}{\partial t} = \nabla \cdot \left[\lambda_{P_2} \nabla \left(\frac{\delta F}{\delta P_2}\right)\right] + k_{on} P \cdot P - k_{off} P_2$$

$$\frac{\partial R}{\partial t} = \nabla \cdot \left[\lambda_R \nabla \left(\frac{\delta F}{\delta R}\right)\right] \qquad F = \int_\Omega \left( \frac{\varepsilon^2}{2}[|\nabla P_2|^2 + |\nabla R|^2] + P_2 \ln P_2 + R \ln R + (1-(P_2+R))\ln(1-(P_2+R)) + \chi_{P_2R} P_2 R + \chi_{P_2S} P_2 (1-(P_2+R)) + \chi_{RS} R (1-(P_2+R)) \right) dx$$

$$P_2 + R + S = 1$$

**C** Model Parameter values

| Parameter | Description | Value |
|---|---|---|
| $\lambda_P$ | Diffusion coefficient of the TDP43 mutant monomer | $1\ \frac{\mu m^2}{sec}$ |
| $\lambda_{P_2}$ | Diffusion coefficient of the TDP43 mutant dimer | $0.5\ \frac{\mu m^2}{sec}$ |
| $\lambda_R$ | Diffusion coefficient of RNA | $0.12\ \frac{\mu m^2}{sec}$ |
| $k_{on}$ | Binding rate of TDP43 monomers to form the TDP43 dimer | $\frac{1}{sec}$ |
| $k_{off}$ | Disassociation rate of the TDP43 dimer | $\frac{0.01}{sec}$ |
| $\chi_{P_2R}$ | Strength of mutant TDP43 dimer and RNA interactions | 4.25 |
| $\chi_{P_2S}$ | Strength of mutant TDP43 dimer and solvent interactions | 4.25 |
| $\chi_{RS}$ | Strength of RNA and solvent interactions | 1 |

**D**

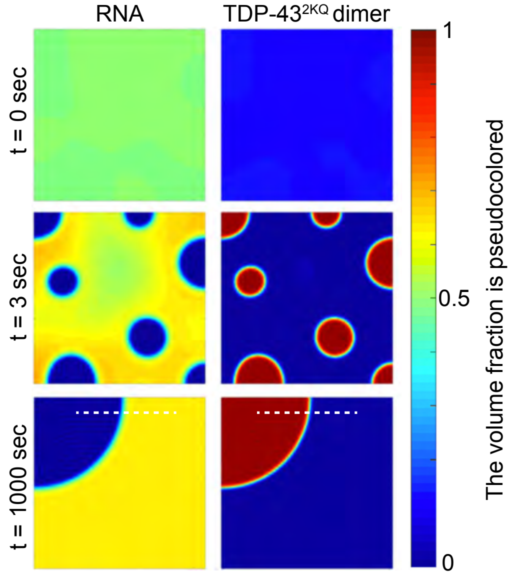


**E**

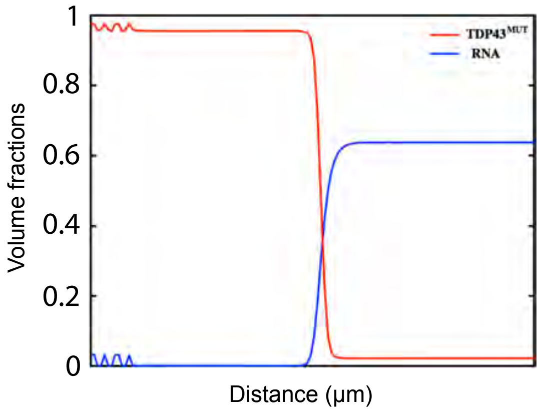


**Fig. S8. A simple model of two component demixing (TDP-43 and RNA) does not support iLSA formation**

**Fig. S8. A simple model of two-component de-mixing (TDP-43 and RNA) does not support iLSA formation.** (**A**) A schematic view describes the modeling of TDP-43$^{2KQ}$ and RNA de-mixing. (**B**) Equations for modeling two component de-mixing. (**C**) Parameters and values. (**D**) The result of modeling. The system starts with a uniform solution of TDP-43$^{2KQ}$ and RNA. Droplets formed and fused at 1~3 seconds, and eventually fused into a big droplet. The color gradient demonstrates relative volume fraction (**E**) Measurement of volume fraction at 1000 second time point for TDP-43 (Red) and RNA (blue).

**A** Equations for Model of demixing including RNA, TDP-43$^{2KQ}$ and "Y"

$$\frac{\partial P}{\partial t} = \nabla \cdot [\lambda_P \nabla P] - 2k_{on} P \cdot P + 2k_{off} P_2$$

$$\frac{\partial P_2}{\partial t} = \nabla \cdot \left[\lambda_{P_2} \nabla \left(\frac{\delta F}{\delta P_2}\right)\right] + k_{on} P \cdot P - k_{off} P_2$$

$$\frac{\partial Y}{\partial t} = \nabla \cdot \left[\lambda_Y \nabla \left(\frac{\delta F}{\delta Y}\right)\right]$$

$$\frac{\partial R}{\partial t} = \nabla \cdot \left[\lambda_R \nabla \left(\frac{\delta F}{\delta R}\right)\right]$$

$$P_2 + R + Y + S = 1$$

$$F = \int_\Omega \left( \begin{array}{c} \frac{\varepsilon^2}{2}\left[|\nabla Y|^2 + |\nabla P_2|^2 + |\nabla R|^2\right] + Y \ln Y + P_2 \ln P_2 + R \ln R \\ +\big(1 - (Y + P_2 + R)\big) \ln\big(1 - (Y + P_2 + R)\big) + \chi_{P_2 Y} P_2 Y + \chi_{P_2 R} P_2 R + \chi_{YR} Y R \\ +\chi_{P_2 S} P_2 \big(1 - (Y + P_2 + R)\big) + \chi_{YS} Y \big(1 - (Y + P_2 + R)\big) + \chi_{RS} R \big(1 - (Y + P_2 + R)\big) \end{array} \right) dx$$

**B** Parameters used in the improved model

| Parameter | Description | Value |
|---|---|---|
| $\lambda_P$ | Diffusion coefficient of the TDP43$^{2KQ}$ monomer | $1 \frac{\mu m^2}{sec}$ |
| $\lambda_{P_2}$ | Diffusion coefficient of the TDP43$^{2KQ}$ dimer | $0.5 \frac{\mu m^2}{sec}$ |
| $\lambda_R$ | Diffusion coefficient of RNA | $0.12 \frac{\mu m^2}{sec}$ |
| $\lambda_Y$ | Diffusion coefficient of Y | $0.5 \frac{\mu m^2}{sec}$ |
| $k_{on}$ | Binding rate of TDP43$^{2KQ}$ monomers to form the dimer | $\frac{1}{sec}$ |
| $k_{off}$ | Disassociation rate of the TDP43$^{2KQ}$ dimer | $\frac{0.01}{sec}$ |
| $\chi_{P_2 Y}$ | Strength of TDP43$^{2KQ}$ dimer and Y interactions | 2.5 |
| $\chi_{P_2 R}$ | Strength of TDP43$^{2KQ}$ dimer and RNA interactions | 4.25 |
| $\chi_{YR}$ | Strength of Y and RNA interactions | 4.25 |
| $\chi_{P_2 S}$ | Strength of TDP43$^{2KQ}$ dimer and solvent interactions | 4.25 |
| $\chi_{YS}$ | Strength of Y a[illegible] solvent interactions | 4.25 |
| $\chi_{RS}$ | Strength of RNA and solvent interactions | 1 |

**Fig. S9. Equations and parameters of three-component de-mixing (TDP-43, RNA, and Y).** (A) Equations for modeling three-component de-mixing. (B) Parameters and values for the modeling. This supplementary figure is in support of Fig. 5A-5C.

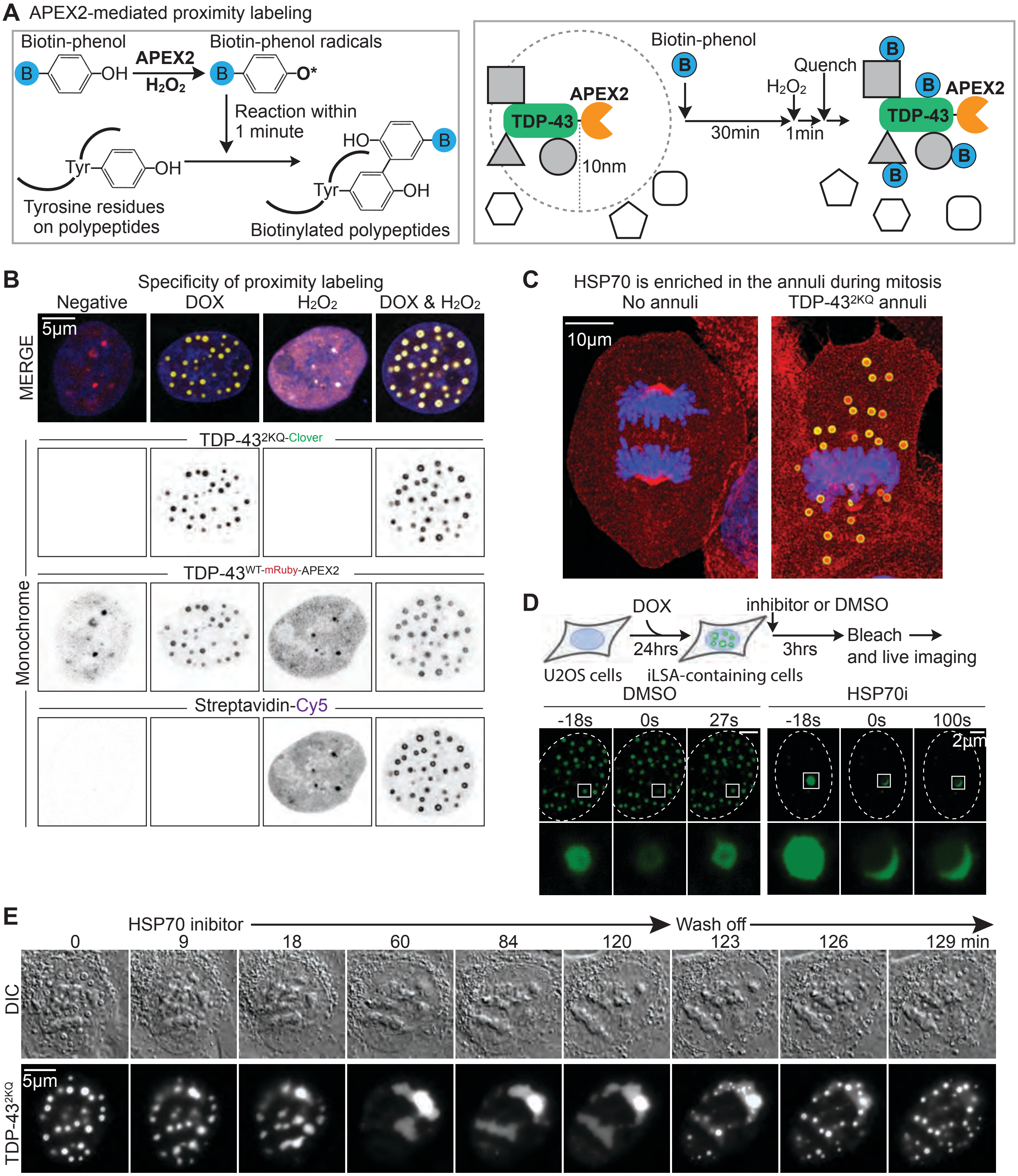


**Fig. S10. HSP70 family proteins were identified as a component enriched inside the iLSA.** (A) A diagram explains the mechanism of proximity labeling. (B) Fluorescent images confirm that iLSA were labeled specifically by proximity labeling. (C) HSP70 proteins stay enriched in the annuli during mitosis. Here fluorescently labeled HSPA6 is shown as an example. (D) FRAP assay shows nuclear TDP-43 gels are not dynamic after HSP70 inhibitor treatment. DMSO was used as vehicle control. (E) A live DIC and fluorescence imaging experiment show reversible annuli to gel formation under HSP70 inhibitor (VER155008). After wash off, the nuclear gel quickly recover to iLSA after 9 minutes.

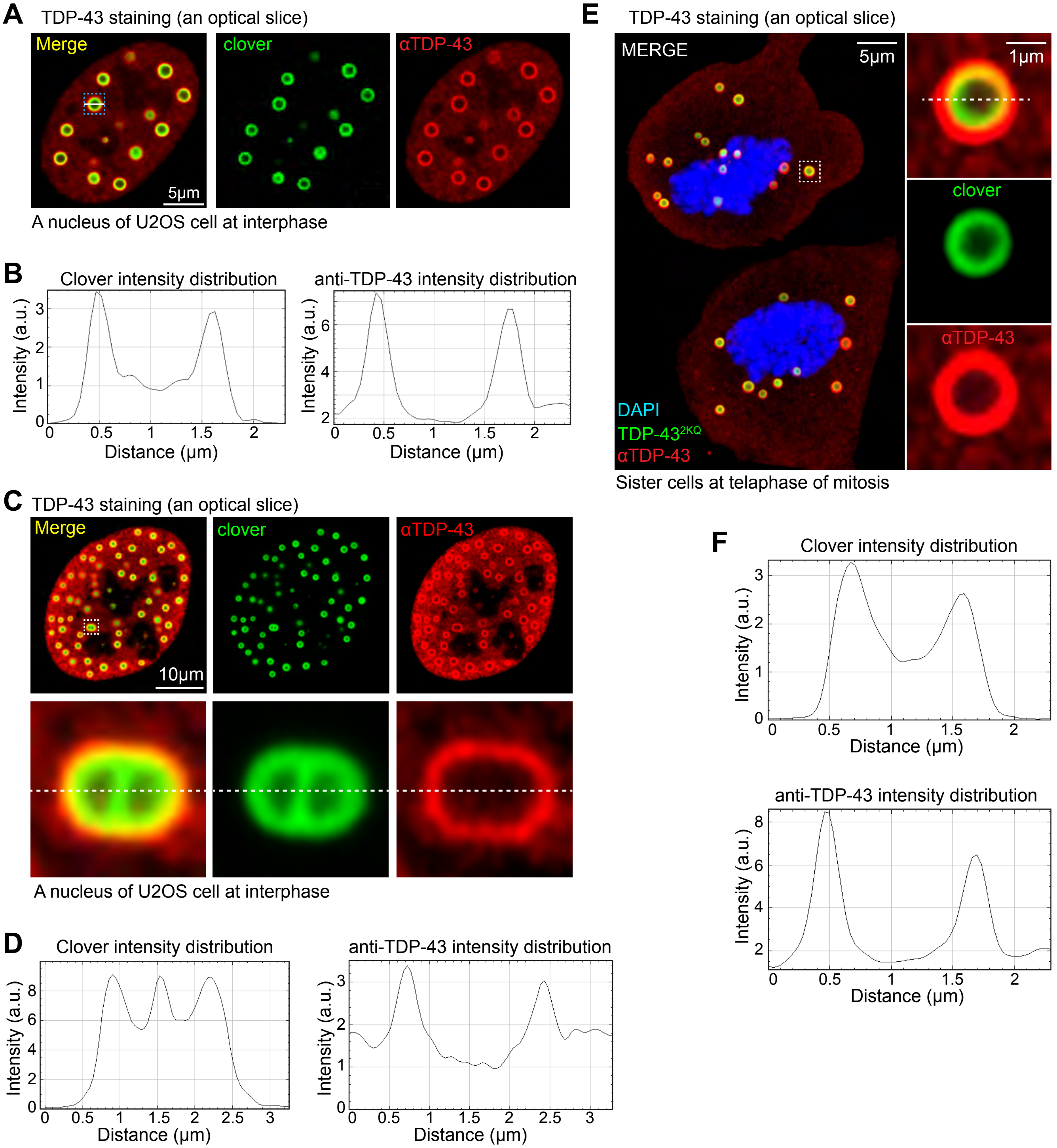


**Fig. S10. Antibodies cannot diffuse through PFA-fixed TDP-43 iLSA. (A) Anti-TDP-43 antibody only stains the outside of the iLSA.** (B) The diameter of the highlighted iLSA measured by clover fluorescence is 1.10 μm, while 1.35 μm by immunostaining. (C) Anti-TDP-43 antibody only stains the outside of a fusion intermediate. Two annuli are in the process of fusion, where the center shows a "bridge" of the same intensity as the edge. However, anti-TDP-43 immunostaining cannot detect the TDP-43 "bridge". This experiment fully demonstrate that fixed annuli are dense meshwork that is a barrier to macromolecules such as antibody. (D) Intensity measurement of the green fluorescence (left) and red fluorescence (right). (E) and (F) TDP-43 annuli at mitosis show similar properties, suggesting the meshwork density remains the same during mitosis.
